# Stabilization of the open conformation of Insulin-Regulated Aminopeptidase by a novel substrate-selective small molecule inhibitor

**DOI:** 10.1101/2024.06.04.597268

**Authors:** Anastasia Mpakali, Galateia Georgaki, Alberto Buson, Alison D. Findlay, Jonathan S. Foot, Francois-Xavier Mauvais, Peter van Endert, Petros Giastas, Dieter W. Hamprecht, Efstratios Stratikos

**Author notes:** Corresponding authors: Dieter Hamprecht, & Efstratios Stratikos, or.

## Abstract

Insulin-Regulated Aminopeptidase (IRAP) is an enzyme with important biological functions and the target of several drug-discovery efforts although no clinically useful inhibitors have been reported yet. We combined *in silico* screening with a medicinal chemistry optimization campaign to discover a nanomolar inhibitor of IRAP based on a pyrazolylpyrimidine scaffold. This compound displays an excellent selectivity profile versus homologous aminopeptidases and kinetic analysis suggests it utilizes an uncompetitive mechanism of action when inhibiting the cleavage of a typical dipeptidic substrate. Surprisingly, the compound is a poor inhibitor of the processing of the physiological cyclic peptide substrate oxytocin and a 10mer antigenic epitope precursor but displays a biphasic inhibition profile for the trimming of a 9mer antigenic peptide and is active in blocking IRAP-dependent cross-presentation of an 8mer epitope. To better understand the mechanism of action and the basis for the unusual substrate selectivity of this inhibitor, we solved the crystal structure of the compound in complex with IRAP. The structure indicated direct zinc(II) engagement by the pyrazolylpyrimidine scaffold and revealed that the compound binds to an open conformation of the enzyme in a pose that should block the conformational transition to the closed conformation previously observed with other low molecular weight inhibitors and hypothesized to be important for catalysis. This compound constitutes the first IRAP inhibitor targeting the active site that utilizes a conformation-specific mechanism of action, provides insight into the intricacies of the IRAP catalytic cycle, and highlights a novel approach to regulating IRAP activity by blocking its conformational rearrangements.

## INTRODUCTION

Insulin-Regulated Aminopeptidase (IRAP) is a zinc metalloprotease that belongs to the oxytocinase subfamily of the M1 family of metalloproteases^1^. IRAP is a transmembrane protein with a large luminal domain that carries its aminopeptidase activity and a small cytoplasmic domain with roles in intracellular trafficking^2,3^. Its enzymatic domain plays several important physiological functions including regulation of immune responses through the generation of antigenic peptides^4^, degradation of placental oxytocin^5^ and the regulation of oxytocin and vasopressin levels in the brain^6^. As a result, several efforts have been initiated to develop pharmacologically useful inhibitors of IRAP’s aminopeptidase activity, in particular in the areas of cognition^7,8^ and fibrosis^9,10^. IRAP inhibitors have been developed either by rational substrate-inspired design or after random chemical or virtual library screening and include pseudophosphinic peptide transition-state-analogues, aryl sulfonamides, benzopyran derivatives, diaminobenzoic acid derivatives, imidazopyridines and cyclic peptide analogues^11–13^. Several of these compounds have been shown to be active in cellular and in *in vivo* systems relating to IRAP biological activities, such as promoting the formation of functional dendritic spines in primary hippocampal neuron cultures^14^, regulation of acetylcholine-mediated vasoconstriction^15^ and glucose tolerance in insulin-resistant Zucker fatty rats^16^. Despite displaying encouraging preclinical efficacy, no clinical applications of IRAP inhibitors have been reported yet, hinting at pharmacological limitations of the compounds developed so far.

The aminopeptidase domain of IRAP consists of four structural domains organized as a concave structure that hosts the catalytic site adjacent to a large cavity^17,18^. Upon ligand binding, IRAP can undergo a conformational change to a more closed configuration in which the internal cavity is sequestered from the external solvent and domain IV makes additional contacts with domains I and II^19,20^. This conformational change is linked to the reconfiguration of structural elements in the catalytic site, including the reorientation of the catalytic Tyr549 and the adjacent GAMEN motif, that result in a presumably more catalytically-optimized active site^19^. As all known crystal structures of IRAP with small molecular weight inhibitors targeting the active site feature this closed conformation, the exact role of the open conformation in the catalytic cycle and its importance in regulating IRAP activity are only now starting to emerge. Indeed, we recently described how one of the most studied inhibitors of IRAP theorized to bind to the active site, is actually allosteric and binds to the open conformation of the enzyme and that this conformation may be enzymatically active for some substrates^21^.

In this study, we further expand on these findings. We utilized virtual screening based on a transition-state analogue inhibitor of IRAP to discover a new series of non-peptidic small molecular weight inhibitors of the enzyme. Medicinal chemistry optimization of an initial hit resulted in a compound with improved potency and excellent selectivity versus homologous enzymes as determined using a standard dipeptidic model substrate. The optimized compound however was found to be a poor inhibitor of cleavage of longer peptides, including the cyclic peptide oxytocin. X-ray crystallographic analysis revealed that the compound binds in the active site but to an open conformation, something not observed before for other active site IRAP inhibitors. This compound features a novel scaffold that targets the IRAP catalytic center, reveals further intricacies of the mechanism of the IRAP catalytic cycle and introduces a novel approach to regulating IRAP’s enzymatic activity. While this reveals potential pitfalls in the search for physiologically relevant inhibitors of this enzyme, it is also suggestive of the prospect of substrate-specific pharmacological modulation of IRAP’s enzymatic activity.

## EXPERIMENTAL METHODS

### Preparation of compound 3

#### Synthesis of 4,6-dichloro-2-(3,5-dimethylpyrazol-1-yl)pyrimidine

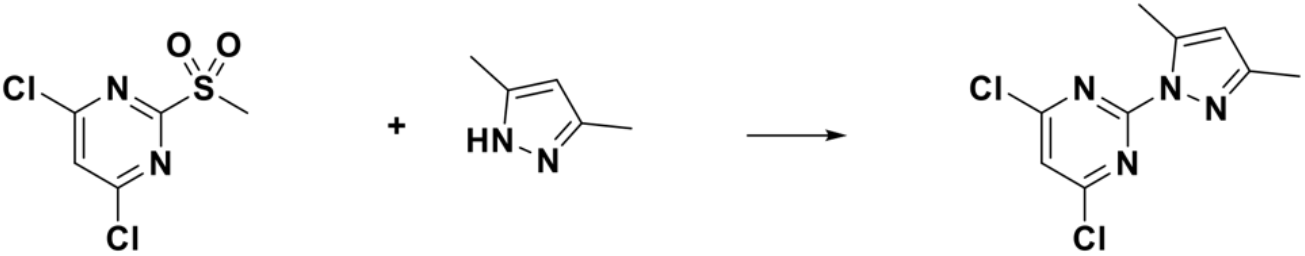

3,5-Dimethylpyrazole (889 mg, 9.25 mmol) was dissolved in CH_2_Cl_2_ (25 mL) and cooled to 0 °C under an atmosphere of nitrogen. Sodium hydride (388 mg, 9.69 mmol) was added portionwise over ∼5 min. The resulting suspension was allowed to stir for 10 min before being added dropwise to a solution of 4,6-dichloro-2-methylsulfonyl-pyrimidine (2.00 g, 8.81 mmol) in CH_2_Cl_2_ (25 mL) at -70 °C. The reaction mixture was stirred at -70 °C for one hour before being quenched via the addition of water (30 mL), allowed to warm to room temperature and transferred to a separatory funnel. The organic layer was washed with water and brine, dried over Na_2_SO_4_ and concentrated *in vacuo*. The crude residue was purified by normal phase flash chromatography (eluting over a gradient of 20-100% EtOAC/hexane) to afford 4,6-dichloro-2-(3,5-dimethylpyrazol-1-yl)pyrimidine (1.58 g, 6.50 mmol, 74%) as a white solid. ^1^H-NMR (300 MHz; CDCl_3_) δ ppm: 7.20 (s, 1H), 6.07 (d, *J* = 1.1 Hz, 1H), 2.66 (d, *J* = 0.9 Hz, 3H), 2.34 (s, 3H).

#### Synthesis of *tert*-butyl 2-[6-chloro-2-(3,5-dimethylpyrazol-1-yl)pyrimidin-4-yl]oxyacetate

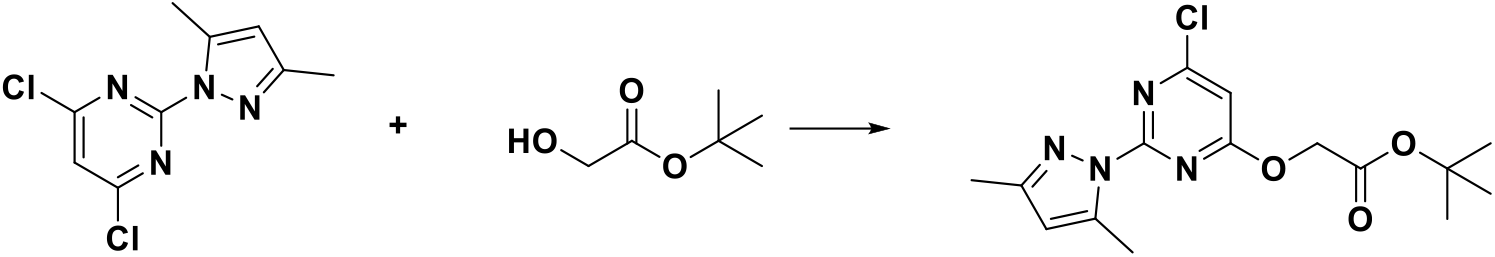

Sodium hexamethyldisilazide (1 M in THF, 7.80 mL, 7.80 mmol) was added dropwise to a solution of *tert*-butyl 2-hydroxyacetate (945 mg, 7.15 mmol) in tetrahydrofuran (15 mL) at 0 °C under an atmosphere of nitrogen and allowed to stir for 5 min. The resulting solution was then added dropwise to a solution of 4,6-dichloro-2-(3,5-dimethylpyrazol-1-yl)pyrimidine (1.58 g, 6.50 mmol) in tetrahydrofuran (15 mL) at 0 °C, and the resultant light brown solution stirred at 0 °C for 1 hour. The reaction was quenched via the addition of water (50 mL) and the resultant mixture extracted with EtOAc (20 mL x 3). The combined organics were washed with brine (30 mL), dried over Na_2_SO_4_ and concentrated *in vacuo* to afford crude *tert*-butyl 2-[6-chloro-2-(3,5-dimethylpyrazol-1-yl)pyrimidin-4-yl]oxyacetate (2.20 g, 5.00 mmol, 77%) as a white solid. ^1^H-NMR (300 MHz; CDCl_3_) δ ppm: δ 6.73 (s, 1H), 6.04 (s, 1H), 4.88 (s, 2H), 2.62 (d, *J* = 0.9 Hz, 3H), 2.33 (s, 3H), 1.46 (s, 9H).

#### Synthesis of 2-[6-chloro-2-(3,5-dimethylpyrazol-1-yl)pyrimidin-4-yl]oxyacetic acid

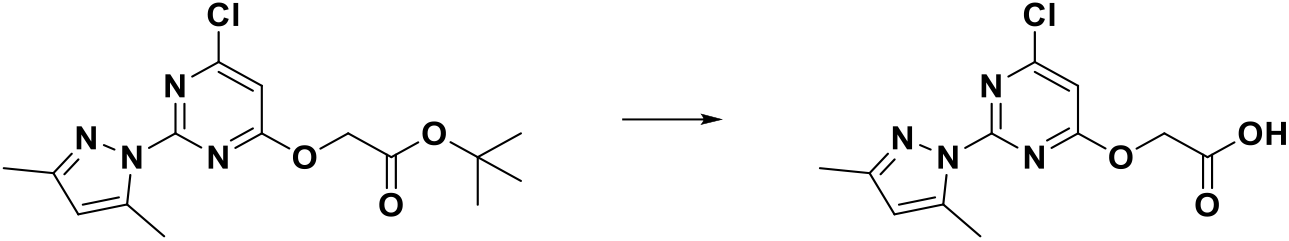

To a stirred solution of *tert*-butyl 2-[6-chloro-2-(3,5-dimethylpyrazol-1-yl)pyrimidin-4-yl]oxyacetate (2.20 g, 5.00 mmol) in CH_2_Cl_2_ (6 mL) was added trifluoroacetic acid (6 mL). The reaction mixture was stirred for 2 hours before concentration *in vacuo* and co-evaporation with CH_2_Cl_2_ (10 mL x 2). The residue was stirred with EtOAc (20 mL) for 30 min. The resulting white solid was collected by filtration, washed the EtOAC (2 mL x 3) and dried under high vacuum to afford 2-[6-chloro-2-(3,5-dimethylpyrazol-1-yl)pyrimidin-4-yl]oxyacetic acid (1.15 g, 4.07 mmol, 81%) as a white solid. 1H NMR (300 MHz, CD_3_OD) δ ppm; 6.94 (s, 1H), 6.17 (tt, J = 0.9, 0.4 Hz, 1H), 5.11 (s, 2H), 2.65 (d, J = 0.9 Hz, 3H), 2.29 (s, 3H).

#### 2-[6-Chloro-2-(3,5-dimethylpyrazol-1-yl)pyrimidin-4-yl]oxy-*N*-methyl-*N*-[(2-methyl-3-pyridyl) methyl]acetamide

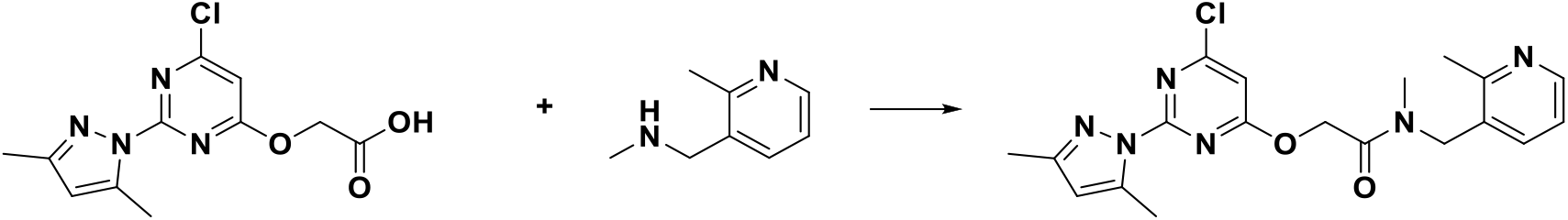

To a stirred solution of of 2-[6-chloro-2-(3,5-dimethylpyrazol-1-yl)pyrimidin-4-yl]oxyacetic acid (350 mg, 1.24 mmol) in CH_2_Cl_2_ (5 mL) at room temperature was added oxalyl chloride (314 µL, 3.71 mmol) dropwise. The reaction mixture was stirred for 30 min before concentration *in vacuo* and co-evaporation with CH_2_Cl_2_ (10 mL x 2). The orange solid so obtained was redissolved in CH_2_Cl_2_ (5 mL) and added dropwise to a mixture of *N*-methyl-1-(2-methylpyridin-3-yl)methanamine (202 mg, 1.49 mmol) and Et_3_N (0.86 mL, 6.19 mmol) in CH_2_Cl_2_ (5 mL) at 0 °C. The resulting solution was stirred at 0 °C for 30 min before being quenched via the addition of water (20 mL) and extracted with CH_2_Cl_2_ (10 mL x 3). The organic layer was dried over Na_2_SO_4_ and concentrated *in vacuo* to afford crude 2-[6-chloro-2-(3,5-dimethylpyrazol-1-yl)pyrimidin-4-yl]oxy-*N*-methyl-*N*-[(2-methyl-3-pyridyl)methyl]acetamide (500 mg, 1.25 mmol, *ca*. 100%) as a brown oil. ^1^H-NMR (300 MHz; CDCl_3_) δ ppm (2:1 mixture of rotamers): 8.35 (dd, J = 4.9, 1.7 Hz, 1H), 7.38 (dd, J = 7.7, 1.6 Hz, 1H), 6.88 (dd, J = 7.7, 4.8 Hz, 1H), 6.77 (s, 1H), 6.04 (d, J = 1.1 Hz, 1H), 5.21 (s, 2H), 4.59 (s, 2H), 3.04 (s, 3H), 2.58 (2s, 3H), 2.47 (s, 3H), 2.27 (s, 3H).

#### 2-[2-(3,5-Dimethylpyrazol-1-yl)-6-(4-methoxyphenyl)pyrimidin-4-yl]oxy-*N*-methyl-*N*-[(2-methyl-3-pyridyl)methyl]acetamide (compound 3)

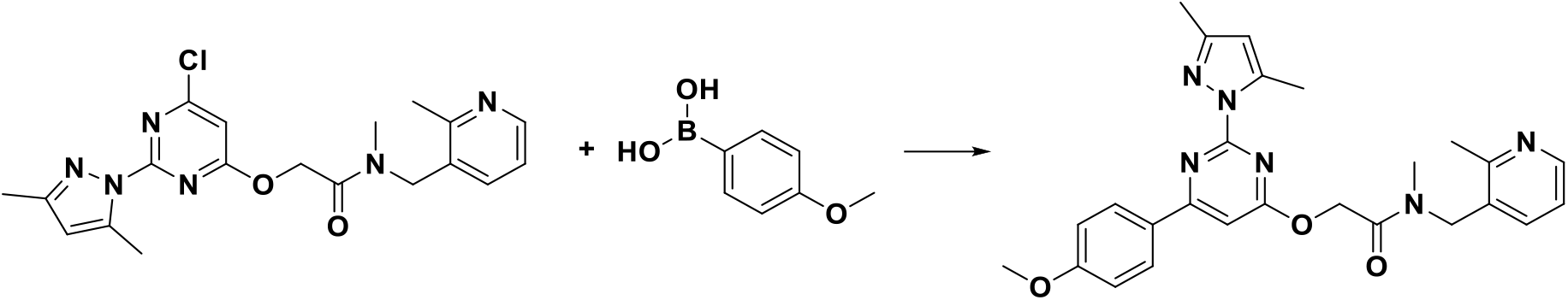

To a stirring solution of 2-[6-chloro-2-(3,5-dimethylpyrazol-1-yl)pyrimidin-4-yl]oxy-*N*-methyl-*N*-[(2-methyl-3-pyridyl)methyl]acetamide (100 mg, 0.25 mmol) in dioxane : water (4 : 1.5 mL) at room temperature was added sodium carbonate (93 mg, 0.87 mmol) and 4-methoxyphenylboronic acid (45 mg, 0.30 mmol). The reaction mixture was degassed with argon, tetrakis(triphenylphosphine)palladium(0) (29 mg, 0.02 mmol) added and heating at 85 °C maintained overnight. The reaction mixture was then diluted with EtOAC (20 mL), dried over Na_2_SO_4_ and adsorbed onto silica gel before purification via normal phase flash chromatography (eluting over a gradient of 5% MeOH in 20-80% EtOAc/CH_2_Cl_2_) to afford 2-[2-(3,5-dimethylpyrazol-1-yl)-6-(4-methoxyphenyl)pyrimidin-4-yl]oxy-*N*-methyl-*N*-[(2-methyl-3-pyridyl)methyl]acetamide, compound **3** (106 mg, 0.22 mmol, 90%) as an off-white foam. ^1^H-NMR (300 MHz; CDCl_3_) δ ppm (mixture of amide rotamers apparent): 8.60 – 8.24 (m, 1H), 8.28 – 7.97 (m, 2H), 7.71 – 7.38 (m, 1H), 7.13 (s, 1H), 7.06 – 6.97 (m, 2H), 6.96 – 6.75 (m, 1H), 6.08 (d, J = 10.5 Hz, 1H), 5.26 (d, J = 34.8 Hz, 2H), 4.64 (s, 2H), 3.89 (s, 3H), 3.10 (d, J = 38.3 Hz, 3H), 2.83 – 2.67 (m, 3H), 2.54 (d, J = 13.4 Hz, 3H), 2.32 (d, J = 10.2 Hz, 3H) ; LRMS (ESI +): *m/z* 473 [M+H]^+^;

See supporting data for preparation of compound 2 and characterization data for compounds 2 and 3.

### Recombinant Enzyme

The production and purification of the soluble extracellular enzymatic domain of IRAP used for enzymatic assays and for crystallization trials has been previously described^18^. Briefly, IRAP was secreted in the medium by stably transfected HEK 293S GnTI^(-)^ cells and purified by affinity chromatography using the Rhodopsin 1D4 tag and the anti-1D4 tag Ab covalently linked to Sepharose beads (Cube Biotech). For crystallization experiments, the protein was additionally purified by size-exclusion chromatography (Superdex 200 16/60 column; GE Healthcare™).

### Enzymatic assays

The activity of the enzyme was estimated by measuring the rate of hydrolysis of the fluorescent substrate L-leucine-7-amido-4-methyl coumarin (Leu-AMC, Sigma–Aldrich, L2145) followed at 460 nm (excitation at 380 nm) using a Spark 10M (TECAN) or a Clariostar (BMG) multimode microplate reader as previously described^22^. For calculation of the *in vitro* IC_50_ values (the transition point at which 50% of the enzymatic activity has been inhibited), experimental data were fitted to a four-parameter [inhibitor] versus response model using Graphpad Prism 8.0. For Michaelis-Menten analysis, initial reaction rates were calculated for each concentration of substrate and the data were fitted to a classical Michaelis-Menten model using Graphpad Prism 8.0.

Oxytocin (Sigma-Aldrich, Cat no. O6379, St. Louis, Missouri, United States) trimming by IRAP was followed by high-performance liquid chromatography as described before^23^. Briefly, 40 μM oxytocin was incubated with 10 nM recombinant IRAP at 37°C for 30 min. The reaction was terminated through acidification by adding 0.25% (v/v) final concentration of trifluoroacetic acid (TFA) and stored at –80°C until analysis. Reactions were analysed using a reversed-phase C18 column (Chromolith™ C-18 column, Merck, Kenilworth, NJ, USA) and a linear gradient (solvent A: 0.05% TFA, 5% acetonitrile in water, solvent B: 0.05% TFA, 50% acetonitrile in water). To estimate fractional digestion, the surface of the peaks corresponding to the substrate and digestion products was integrated and compared to a control reaction. To estimate IC_50_ values, reactions were performed with increasing concentrations of compound and data were fitted to a variable slope log(inhibitor) versus response model using GraphPad Prism 8.0. For the Michaelis-Menten analysis, the same assay was performed using increasing concentrations of substrate in the absence or presence of 100 μM compound **3**. Linear peptide trimming of the extended antigenic peptide precursor LDRASFIKNL (JPT Peptide Technologies GmbH, derived from the epitope DRASFIKNL from Collagen alpha-2(VI) chain^24^) and antigenic peptide YTAFTIPSI (derived from the Gag-Pol polyprotein of human immunodeficiency virus^25^) was performed similarly to oxytocin trimming. For these reactions, 30 μM of the peptide was mixed with 10 or 20 nM recombinant IRAP in 50 mM HEPES, pH 7.0, 150 mM NaCl and the mixture incubated for 45 min at 37°C.

### Crystallization and data collection

Crystals of IRAP complexes with ligand were obtained after screening using the commercially available PACT-premier crystallization screen (Molecular Dimensions™). Crystals were grown at 16 °C by the sitting-drop vapor-diffusion method in 0.1M SPG buffer (Succinic acid, Phosphate, Glycine), pH 9.0 and 25% (w/v) polyethylene glycol (PEG) (A6 condition PACT). One hour before crystallization tests, purified IRAP at 9.8 mg/mL was mixed with the compound at a molar ratio of 1:5 and the mixture was incubated at 4 °C. Cryoprotection was performed by rapid immersion in a solution containing the precipitant and 20% ethylene glycol for 5-10 seconds and then crystals were flash-frozen in liquid N_2_. The frozen crystals were shipped in a Taylor-Wharton CX100 dry shipper to the Synchrotron facility. Diffraction datasets were collected at 100 K at a wavelength of 0.976 Å on beamline P13 of Petra III, EMBL, Hamburg, Germany. The reflections were integrated and scaled with XDS and the space group was determined with POINTLESS. The best dataset for compound **3** displayed diffraction to 3.3 Å and space group P 1 21 1 with a=68.229 Å, b=255.121Å, c=73.274 Å, and β=110 °. The resolution limit was set at 3.5 Å to ensure that the CC1/2 value in the outer shell is higher than 40%, the mean(I/σ) is higher than 1.0, and the completeness is approximately 100% in all resolution shells.

### Structure determination and refinement

The crystal structure was solved by molecular replacement with PHASER using as a search model the structure of IRAP with PDB ID: 4Z7I which corresponds to the open conformation of the enzyme. No reliable solutions were found when using models that correspond to the closed conformation of IRAP (PDB ID: 5MJ6). Two protein molecules were found in the asymmetric unit (referred to as chains A and B). Electron density that was not attributable to the protein molecule was found in each set adjacent to the active site and interpreted to belong to the bound ligand. Structure refinement was conducted with PHENIX using restrained refinement and non-crystallographic symmetry restraints. Model building and real space refinement were performed in Coot. The structure of IRAP in complex with compound **3** converged to R and R_free_ of 24.00 and 27.67 %, respectively. No density was visible before residue 160 and between residues 639-647 in chain A, while in chain B, no density was visible before residue 161 and between residues 222-226, 228 and 639-646. The model also includes 32 molecules of N-acetyl-D-glucosamine, 6 molecules beta-D-mannose, and 2 molecules alpha-D-mannopyranose.

### *In vitro* antigen cross-presentation

MutuDC cells (an immortalized C57BL/6 murine splenic conventional dendritic cell type 1 (cDC1)-like line (a gift from Dr. Hans Acha-Orbea^26^) were plated into a 96-well culture plate (30,000 cells/well) with round bottoms (Nunc) in culture medium (RPMI-1640 medium (Merck) + 10% heat-inactivated Fetal Bovine Serum (Eurobio) + 2 mM L-Glutamine (Merck) + 1% Penicillin/Streptomycin, 50 μM β-mercaptoethanol (Gibco) + 25 mM HEPES (Thermo Fisher Scientific) + 1X Non-Essential amino acids (Gibco), 1 mM Sodium Pyruvate (Thermo Fisher Scientific)), 5% CO2, 37°C. Compound stocks were prepared at 10 mM in DMSO and diluted in serum-free medium at the appropriate concentrations. 30 min after plating the cells, the medium was removed and serial dilutions of compound or DMSO (vehicle, Merck) were added to the wells and incubated for 30-40 min. Ovalbumin (OVA, Worthington) or an irrelevant antigen, Bovine Serum Albumin (BSA, Merck), were then provided to the cells at a concentration of 0.5 mg/mL for 16-20 hours, in the presence of the inhibitor or vehicle (DMSO). As a positive control, serial dilutions of synthetic SIINFEKL peptide were provided to the cells in the presence of compound (30 μmol/L) or vehicle. At the end of incubation, cells were washed three times in PBS before the addition of B3Z hybridoma (a gift from N. Shastri^27^) in culture medium for another 24h (ratio 1:1). Culture supernatants were frozen at -20°C for up to 10 days before dosing IL-2 by a sandwich ELISA technique. In brief, Nunc Maxisorp ELISA plates (Nunc) were coated overnight with purified anti-IL2 antibody (clone JES6-1A12, Biolegend, 2 μg/mL) diluted in PBS (50 μL/well). Coating solution was discarded, and plates were blocked with a PBS 10% Fetal Bovine Serum (200 μL/well) for 30 min at room temperature. After discarding blocking solution, supernatants or serial dilutions of mouse IL2 recombinant protein (Peprotech, ThermoFisher Scientific) (100 μL/well) were added for 60-90 min to the plates at room temperature, with gentle shaking (300 rpm). Plates were then washed twice with PBS Tween20 0.05% (300 μL/well), and biotin-conjugated anti-IL2 antibody (clone JES6-5H4, Biolegend, 2 μg/mL) diluted in PBS (50 μL/well) was incubated for 45 min at room temperature with gentle shaking (300 rpm). After 5 washes in PBS Tween20 0.05% (300 μL/well), HRP-streptavidin (BD Biosciences, 1:2000) diluted in PBS was added for 30 min (500 μL/well) at room temperature with gentle shaking (300 rpm). After 5 washes in PBS Tween20 0.05% (300 μL/well), TMB substrate (Cell Signaling Technology) reagent was added (100 μL/well) for 5-10 min at room temperature to reveal bound cytokine and reaction was blocked by adding HCl 2N (Merck) solution (20 μL/well). The absorbance at 450 nm was read using a Mithras LB940 reader (Berthold Technologies). Each condition was performed in duplicate. IL-2 OD readings for OVA-incubated cells were corrected for background values obtained with BSA-incubated cells, before their conversion into IL-2 concentrations based on the linear curve of mouse recombinant IL-2 standards. The results are presented as the means of 4 independent experiments. Statistical analysis was performed using RStudio (version 2022.12.0+353, with R version 4.2.2). In brief, statistical significance was determined using one-way ANOVA test and post-hoc pairwise comparisons using t tests with pooled SD and the Benjamini-Hochberg for P-value adjustment method.

## RESULTS

*In silico* screening was carried out using the BLAZE^TM^ advanced ligand-based virtual screening suite (Cresset) starting from the crystal structure of IRAP in complex with DG025 transition-state analogue (PDB: 4Z7I), and using a truncated form (first 4 residues) of the inhibitor. Screening was carried out on a database of commercially available chemicals and also on the CHEMBL database. Filters were then applied to remove reactive groups, pan-assay interference compounds (PAINs) and known zinc-binding motifs, and the hits thus obtained separated into structural classes/clusters. Top-scoring hits from each cluster that were commercially available through ChemBridge^TM^ were purchased and screened initially at 10 μM for inhibition *in vitro*. Those showing >20% inhibition were further progressed to obtain IC_50_ values.

These efforts led to the discovery of compound **1**, which displayed modest potency (IC_50_ = 55 µM) against recombinant human IRAP. A medicinal chemistry multi-parameter optimization campaign was then undertaken, primarily to increase potency against the target and improve the physiochemical properties of the initial hit. Modification of the template and northern substituent regions of the initial hit (compound **1**) led to the discovery of the significantly more potent compound **2** (IC_50_= 4 μM) which was optimized further, resulting in compound **3** which combines a further increase in potency with a balanced distribution of polar features.

**Scheme 1:**
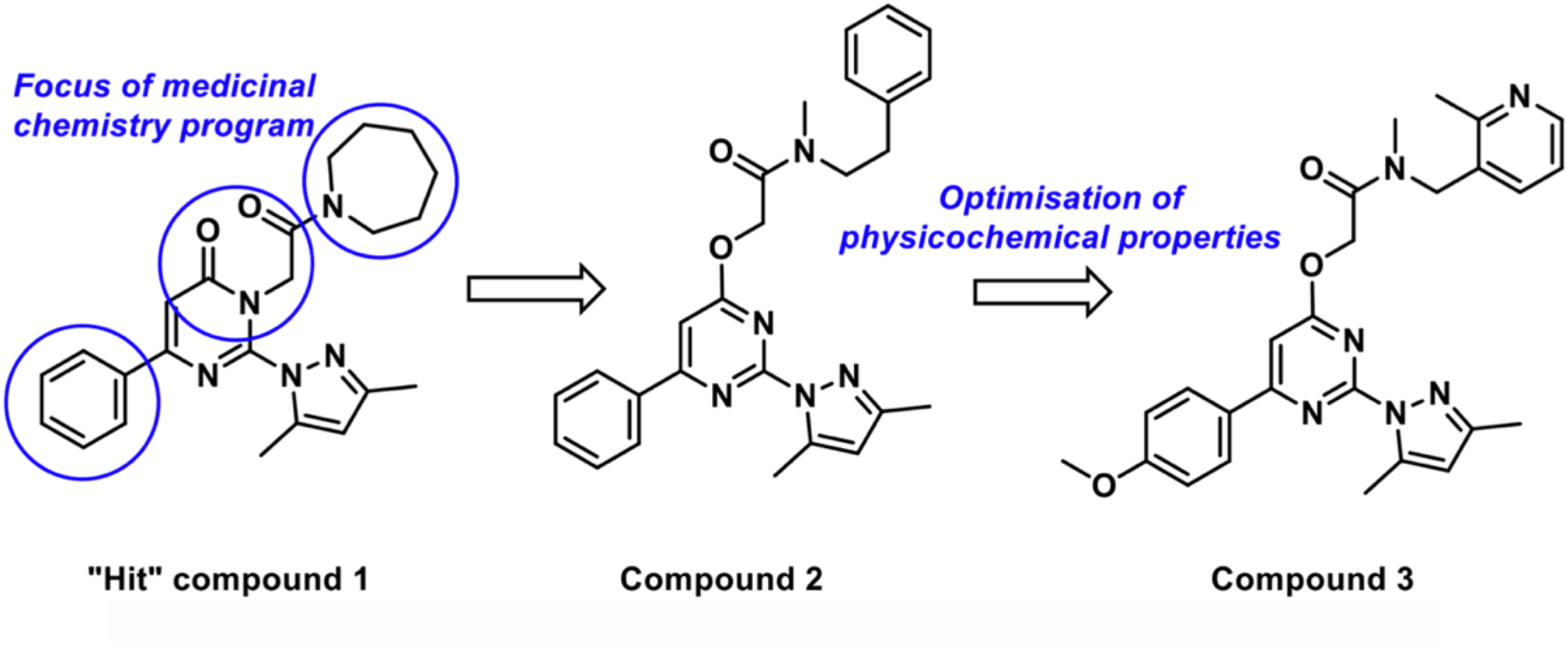
Chemical structures of initial hit compound **1**, intermediately optimized compound **2** and resulting compound **3**, after further optimizations.

Using the fluorogenic substrate Leu-AMC to determine inhibitor potency, compound **3** displayed an IC_50_ of 157 nM against recombinant IRAP and very high selectivity (>200 fold) over the homologous enzymes Endoplasmic Reticulum Aminopeptidase 1 (ERAP1) and 2 (ERAP2) (no inhibitory activity detected up to 30 μM) (Figure 1A). Michaelis-Menten analysis suggested that the inhibitor utilizes an uncompetitive mechanism by decreasing both the V_max_ and the K_M_ of the enzyme (Figure 1B, C). This mode of inhibition was surprising given that compound **3** has been derived from hits of a virtual screen based on direct binders at the catalytic site of IRAP.

**Figure 1:**
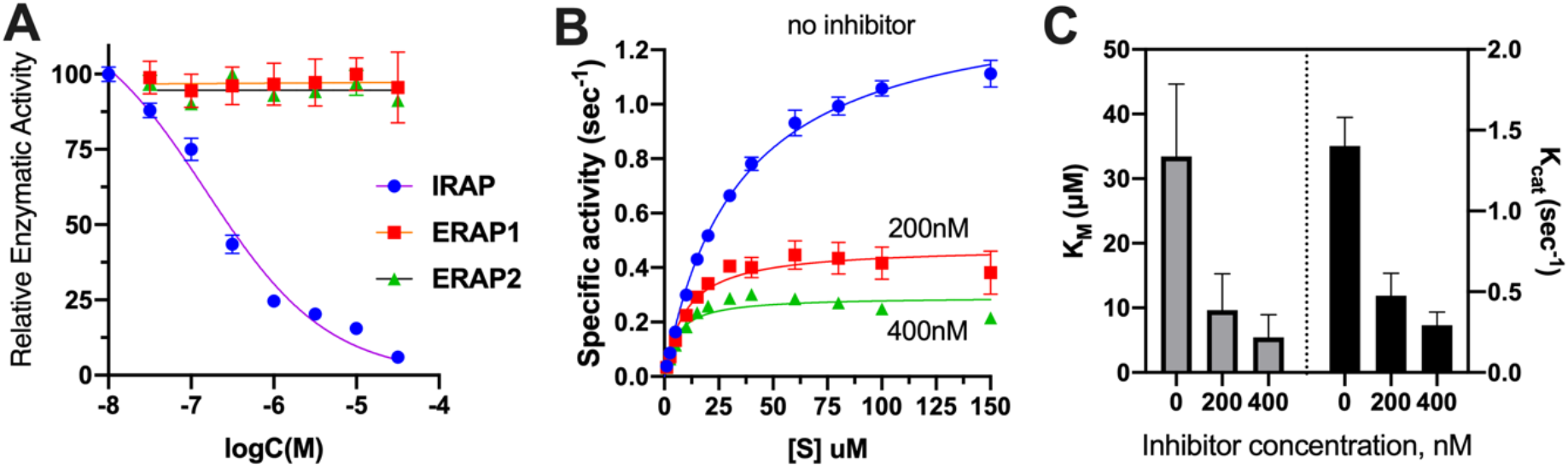
**Panel A**, titration of compound **3** while following the enzymatic activity of IRAP and homologous enzymes ERAP1 and ERAP2 using the Leu-AMC fluorogenic substrate. **Panel B**, Michaelis-Menten analysis of Leu-AMC substrate digestion by IRAP in the presence or absence of compound **3. Panel C**, calculated parameters k_cat_ and K_M_ based on the data shown in Panel B. Error bars are calculated from independent replicates (n=3).

IRAP can degrade and inactivate cyclic peptide hormones such as oxytocin and vasopressin and through this action plays roles physiologically, such as in cognitive functions^28^. It can also trim linear precursors of antigenic peptides or mature antigenic peptides as part of its role in the adaptive immune response and the pathway of cross-presentation^4^. Although the mechanism of processing of macrocyclic peptides by IRAP is not fully understood, the flexibility of the GAMEN loop has been proposed to be important^20^. We thus investigated substrate-dependent potency of inhibition by compound **3** versus three other types of IRAP substrates, a cyclic bioactive peptide, a linear antigenic epitope precursor and a model antigenic peptide. We utilized a previously developed HPLC-based assay for following the trimming of peptides by IRAP^23^ (representative chromatograms for the degradation of oxytocin are shown in Figure 2A). Using this assay, we discovered that compound **3** was a much weaker inhibitor for the cyclic substrate oxytocin with estimated IC_50_ values over 100 μM (an accurate IC_50_ could not be determined due to limited solubility of the compound at high concentrations) (Figure 2B). We obtained similar weak inhibition results when using a 10mer model precursor antigenic epitope (Figure 2C). In contrast, the transition-state analogue DG026 which has been shown to bind to the active site of the closed conformation, was able to fully inhibit oxytocin trimming at 1 μM (Supporting Figure 1)^19^. When following the degradation of a 9mer antigenic epitope, compound **3** exhibited a biphasic behavior with partial inhibition with an IC_50_ of about 100 nM and a second phase with a much weaker ∼100 μM IC_50_ (Figure 2D). These results suggest that the efficacy of compound **3** is dependent on the nature of the substrate and is possibly affected by the substrate length. Remarkably, this profile is similar to a previously characterized allosteric inhibitor of IRAP, which also displayed an uncompetitive mechanism of action towards the Leu-AMC substrate.^21^

**Figure 2:**
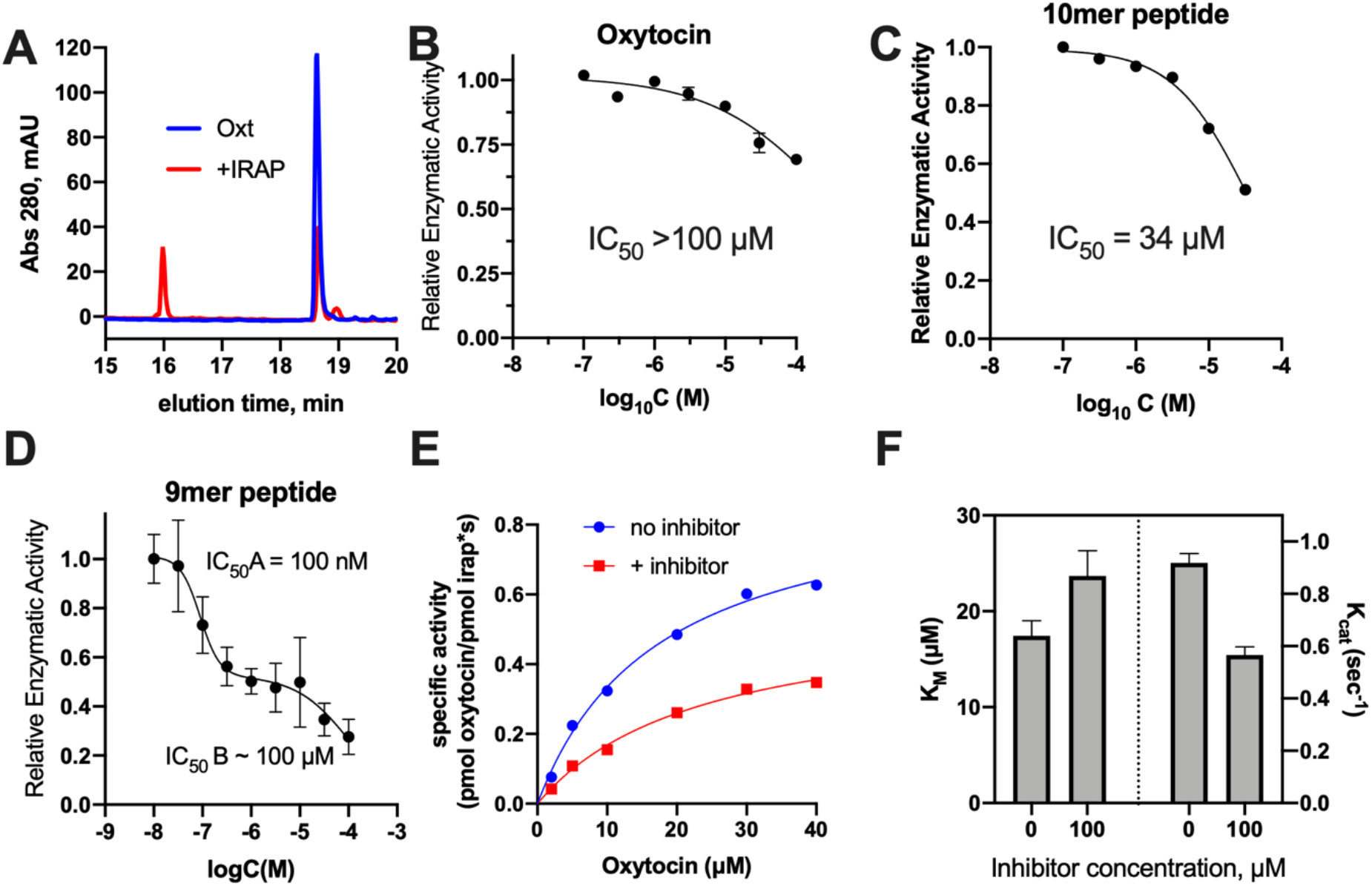
**Panel A**, representative chromatograms of the oxytocin trimming assay: Reversed-phase HPLC chromatograms of oxytocin control and oxytocin incubated with 10 nM IRAP for 30 min at 37 °C. **Panel B**, titration of compound **3** inhibits trimming of oxytocin, but with a very weak IC_50_ estimated to be over 100 μM. **Panel C**. titration of compound **3** inhibits trimming of the 10mer linear antigenic peptide precursor with the sequence LDRASFIKNL, but with a very weak IC_50_ estimated to be 34 μM. **Panel D**, titration of compound **3** inhibits trimming of the 9mer linear antigenic epitope with the sequence YTAFTIPSI. Data for panels B, C were fitted to a variable slope log(inhibitor) versus response model and data for panel D were fitted to a biphasic dose-response curve using GraphPad Prism 8.0. **Panel E**, Michaelis-Menten analysis of oxytocin digestion by IRAP in the presence or absence of 100 μM compound **3. Panel F**, calculated parameters k_cat_ and K_M_ based on the data shown in Panel E. Error bars are calculated from independent replicates and only shown if are significantly wider than the size of the data point (n=2 for panels B,C,E and F; n=4 for panel D).

To better understand this phenomenon, we performed a Michaelis-Menten analysis with compound **3** for the oxytocin substrate (Figure 2E). The inhibitor acted by a mixed non-competitive mechanism, decreasing the *k*_cat_ and increasing the K_M_ parameter (Figure 2F). A typical interpretation of this result is that compound **3** acts by binding to an allosteric site. However, this again seems surprising since compound **3** has been identified and optimized based on ligands that bind to the active site of the enzyme.

Given the complex substrate-selectivity of compound **3**, we tested its efficacy using a cell-based assay based on the presentation of the epitope SL8 (sequence: SIINFEKL), derived through ovalbumin degradation, by an immortalized C57BL/6 murine splenic conventional dendritic cell, type 1-like line, MutuDC cells as described in the methods section. The addition of synthetic peptide SL8 to MuTu DCs, resulted in direct presentation and was used as a positive control. Compound **3** had no apparent effect on direct presentation, which was expected since this process is IRAP-independent (Figure 3A)^4^. The SL8 epitope can be presented through cross-presentation when the ovalbumin is added to the media and internalized by the cells. In this case, presentation was reduced in a dose-dependent fashion by the addition of compound **3** (Figure 7B) to about 40% of the initial signal. Although this limited effect may be related to the inability of the compound to fully inhibit IRAP, it is more likely due to the involvement of ERAP1^29^ or also ERAP2 in the process, two enzymes not inhibited by compound **3**. To quantitate the effect, we fit the data shown in Figure 3B to a simple inhibition model, which allowed the calculation of an EC_50_ value of 640 nM, which is only 4-fold less than the *in vitro* potency of compound **3**, suggesting that the compound is efficient in blocking the generation of the 8mer SL8 peptide from ovalbumin in mouse cells (Figure 3C).

**Figure 3:**
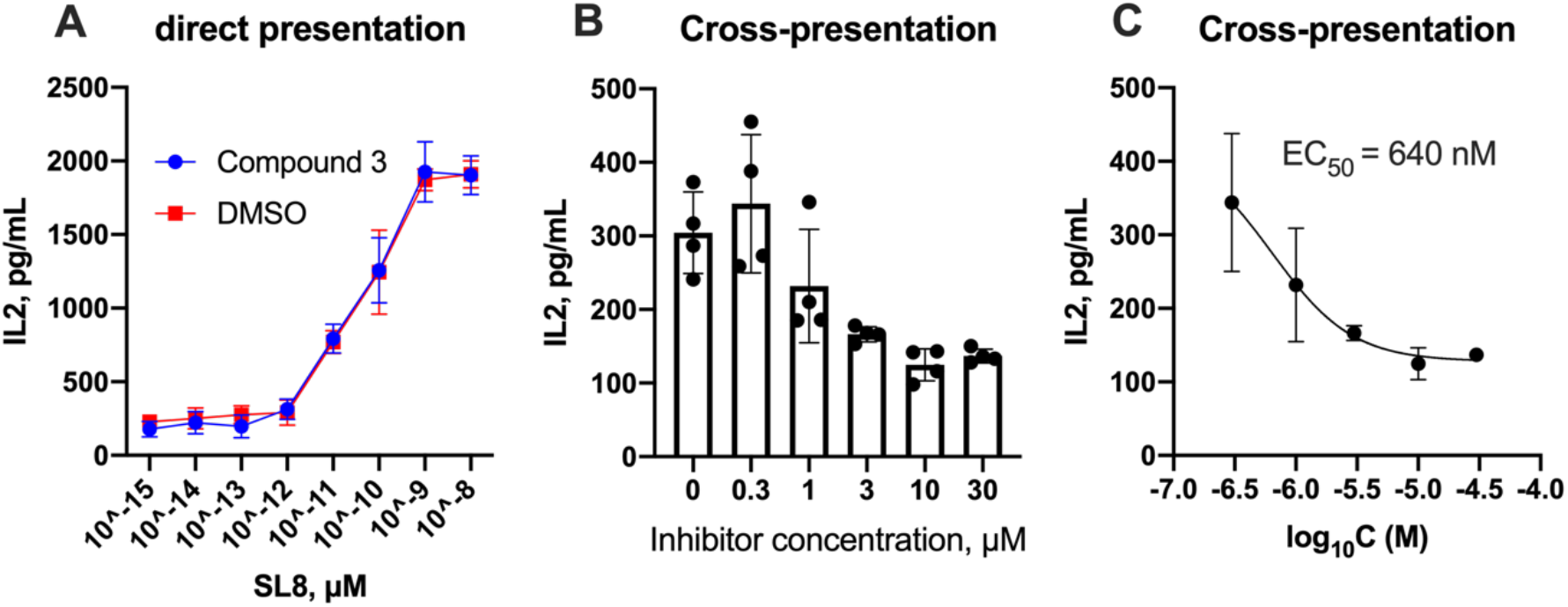
Evaluation of the effect of compound **3** on the SL8 ovalbumin epitope presentation by MutuDC cells. **Panel A**, quantitation of direct presentation after epitope addition. The dose-dependent effect of titrating peptide SL8 is not affected by the compound. **Panel B**, cross-presentation of the SL8 epitope after the addition of ovalbumin. Compound **3** reduces cross-presentation in a dose-dependent manner. **Panel C**, data from panel B were fit to a dose-dependent inhibition model to allow for the calculation of EC_50_.

To shed light onto the mechanism of action of this inhibitor in structural terms, we co-crystallized compound **3** with the extracellular domain of IRAP that holds the enzymatic activity and solved the crystal structure at 3.5 Å. Despite the overall low resolution of the structure, we clearly detected sufficient additional electron density at the active site adjacent to the catalytic zinc(II) atom (Figure 4A). This density was used to build and refine a model of the bound inhibitor as shown in Figure 4B. Crystallographic data collection and refinement statistics are shown in Table 1. The inhibitor was found to engage the active site zinc(II) atom in a bidentate fashion utilizing the pyrazole and pyrimidine groups (Figure 4C). The methoxybenzene group and the methylpyridine group are accommodated into two pockets (indicated as P1 and P2 in Figure 4D) that correspond to specificity pockets of the enzyme. Overall, the inhibitor makes atomic contacts (within 4 Å) with at least 12 residue sidechains in the active site suggesting a highly optimized interaction (Figure 4C). These atomic interactions include T-shaped π−π interaction with Phe544 a residue shown to be important for the specificity of the enzyme^30^.

**Table 1:**
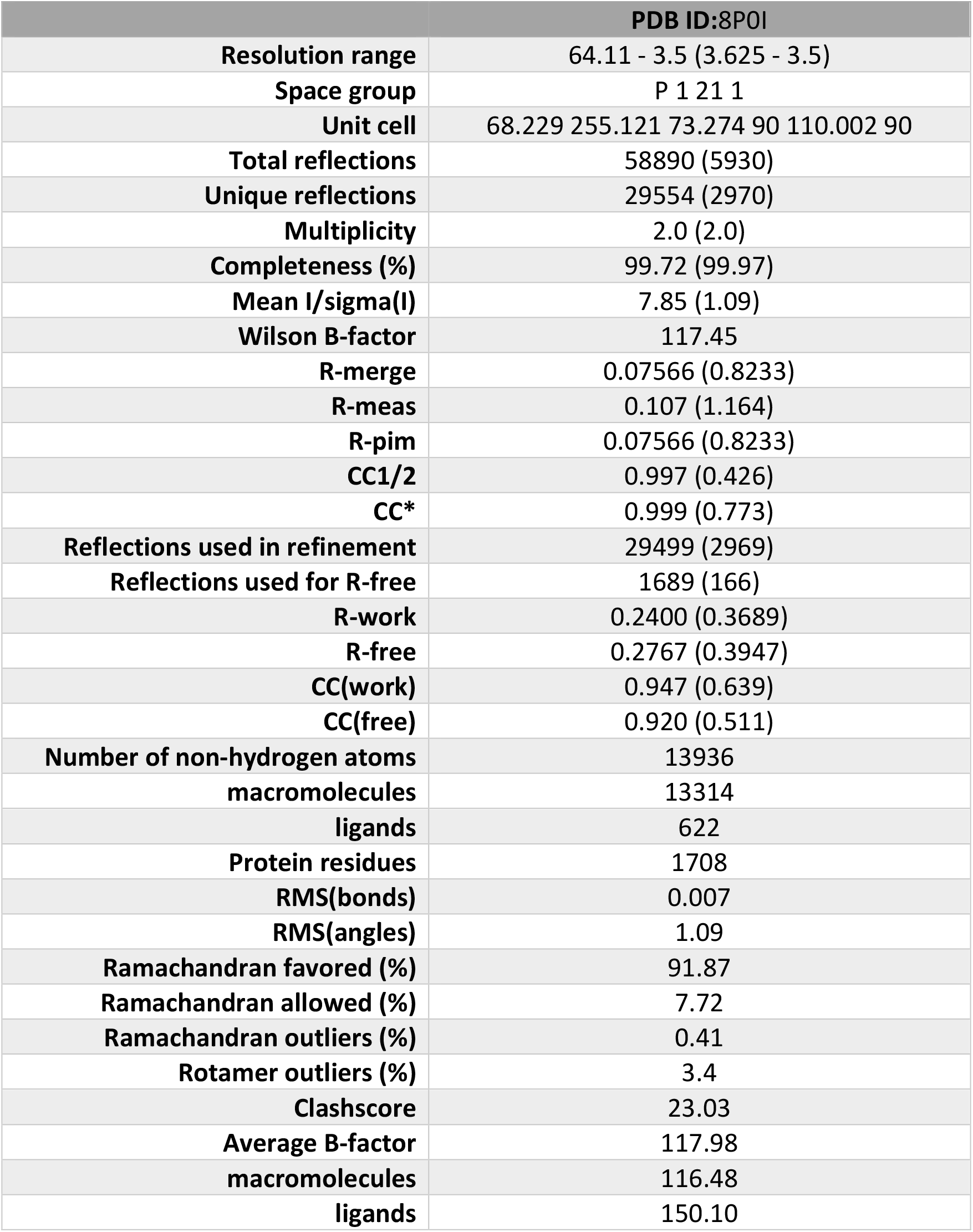
Data collection and refinement statistics. Statistics for the highest-resolution shell are shown in parentheses.

**Figure 4:**
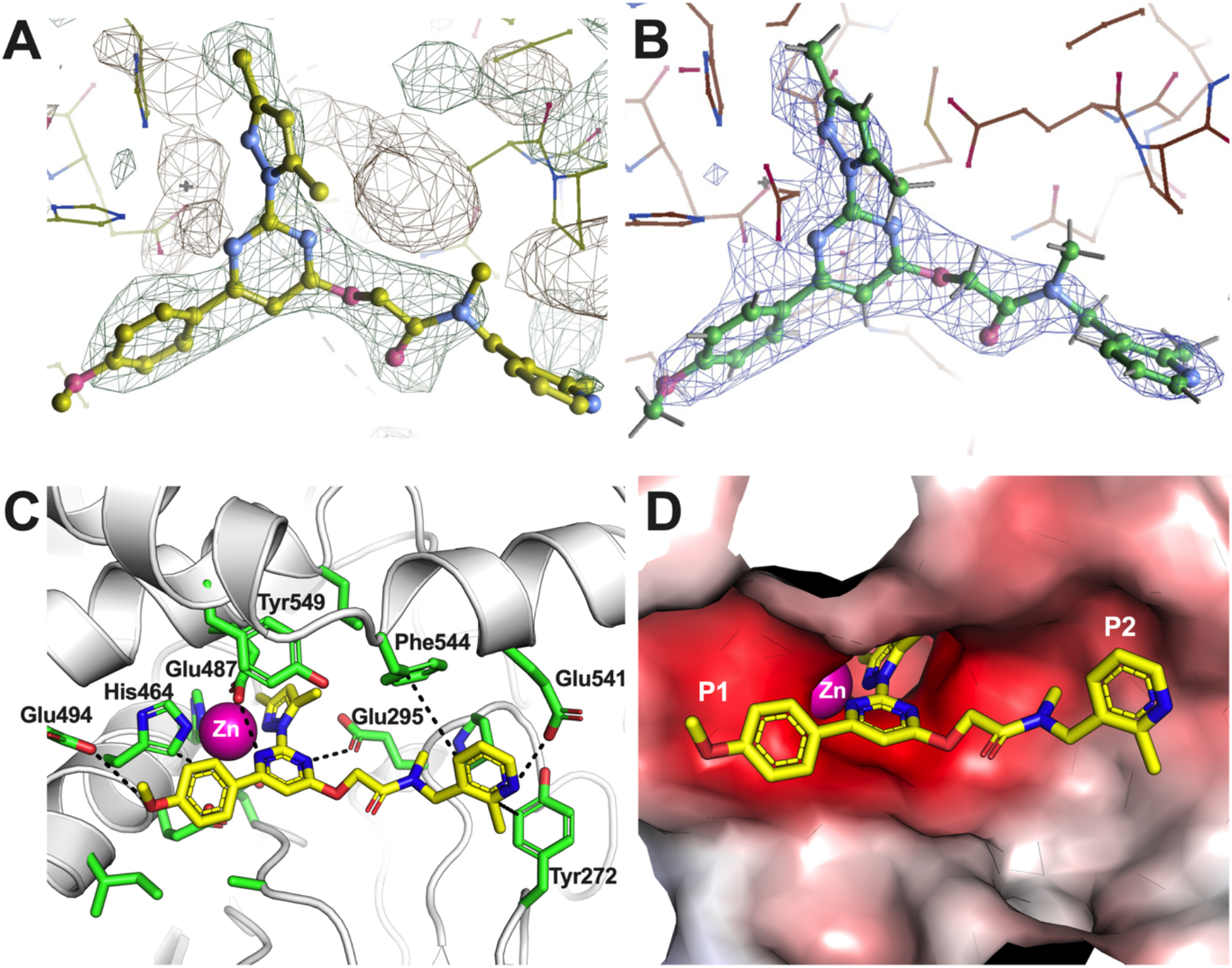
**Panel A**, F_o_-F_c_ difference map showing the density (in mesh representation) in the active site of chain A where the ligand was modeled (shown in ball and stick representation). Sigma was set at 2.0. **Panel B**, 2F_o_ − F_c_ electron density map (blue mesh) contoured at 1.0 sigma after refinement. Ligand is shown in ball and stick representation. **Panel C**, schematic representation of compound **3** in complex with IRAP. Compound is shown in stick representation colored by atom (carbon = yellow, oxygen = red, nitrogen = blue). IRAP sidechains that come within 4 Å of the inhibitor are shown in stick representation. The overall structure of IRAP is shown in white cartoon representation. Dotted lines indicated putative atomic interactions that stabilize the inhibitor. The catalytic Zn(II) atom is shown as a magenta sphere. **Panel D**, compound **3** in the active site of IRAP. Compound is colored as in panel C. IRAP is depicted in surface representation colored by local electrostatic potential (red=negative, blue=positive, white=hydrophobic). P1 and P2 pockets are indicated.

The overall conformation of the enzyme is distinct from all previously published structures determined in complex with active site binding small-molecule inhibitors and corresponds to the open conformation in which domain IV moves away from domains I and II using domain III as a hinge (Figure 5). The observed conformation is consistent with the original template used for the virtual screening which corresponds to the open conformation of IRAP but comes in sharp contrast with all other known structures of IRAP with small inhibitors occupying the active site (PDB IDs: 6YDX,7ZYF, 5MJ6)^19,20,31^, suggesting that the previously stipulated hypothesis that active-site binding induces conformational closing of the enzyme may not hold true for all ligands^19,20^.

**Figure 5:**
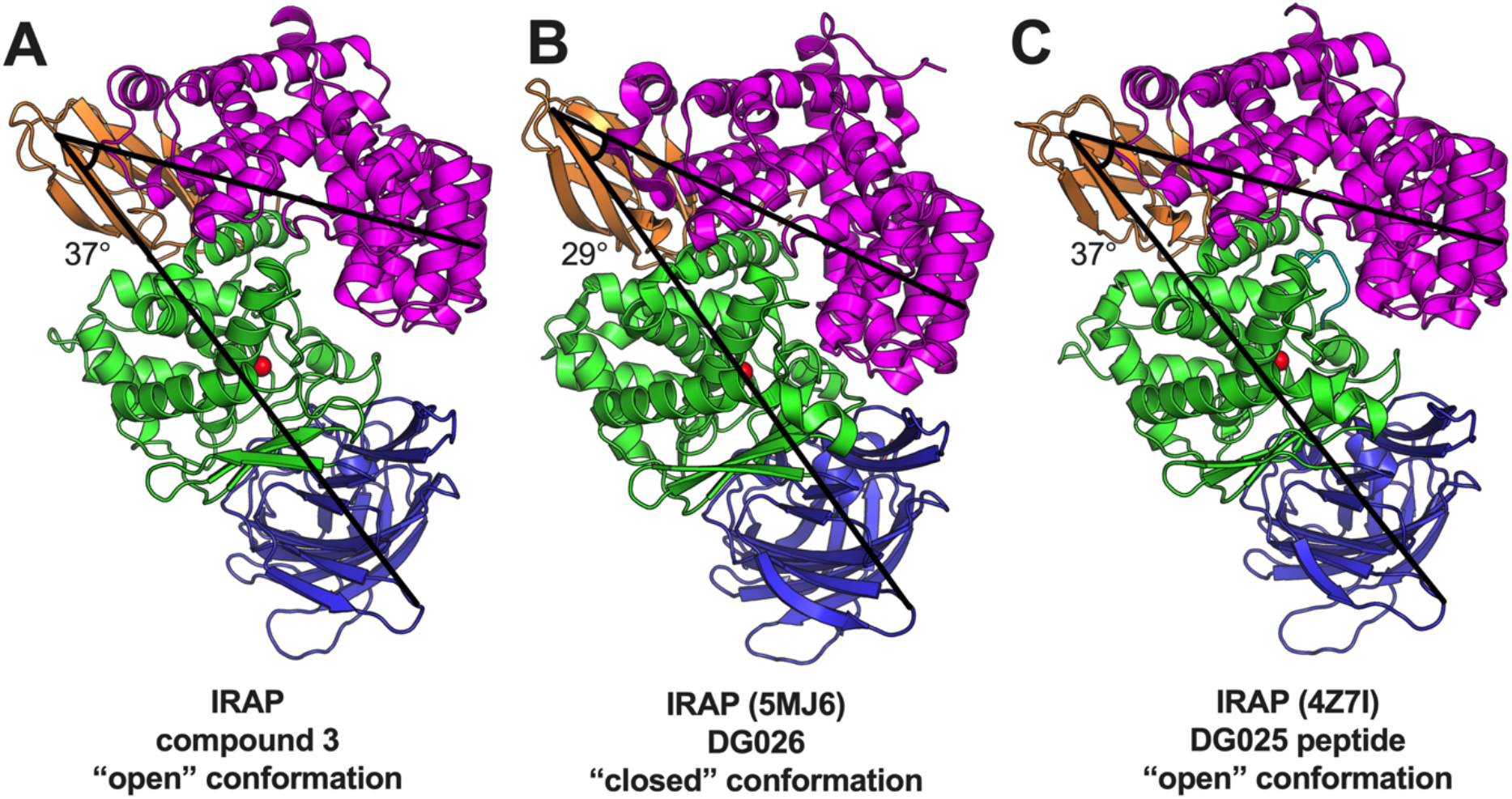
Cartoon representation of the overall domain organization of IRAP in complex with compound **3** (**Panel A**), transition-state analogue DG026 (**Panel B**, PDB ID 5MJ6) and 10mer peptidic analogue DG025 (**Panel C**, PDB ID 4Z7I). Domain I is colored blue, domain II green, domain III orange and domain IV magenta. Active site zinc(II) atom is shown as a red sphere. To help visualize the differences in relative domain orientation, lines between Glu661-Asp341 and Glu661-Lys911 are drawn and the angle between them is indicated.

To gain insight on why compound **3** was found bound onto the open conformation of IRAP in contrast with all other active-site binding small-molecule compounds analyzed before, we aligned the IRAP/compound **3** complex with the crystallographic structure of IRAP in complex with a transition-state analogue which corresponds to the closed conformation of the enzyme (Figure 6A). Visual inspection of the aligned structures suggested that observed steric clashes can explain why the binding pose of compound **3** is incompatible with the closed conformation of IRAP. Specifically, several active site residues reposition during closing at positions that would clash with compound **3** binding, including Tyr549, Phe544 and Glu487. Most importantly, the major motion of domain IV of IRAP would bring Ser960, a residue at the turn between two domain IV helices, in direct steric clash with the 2-methylpyridine moiety of the inhibitor. Thus, it appears that the structure of compound **3** is optimized for the open conformation of IRAP and that binding to the active site in the open conformation blocks conformational closing. Interestingly, the allosteric inhibitor HFI-419^21^ binds in the open conformation of IRAP at the same general location where the 2-methylpyridine group of compound **3** is located, suggesting the occupation of that site may be key for blocking the conformational transition to the closed conformation (Figure 6B).

**Figure 6:**
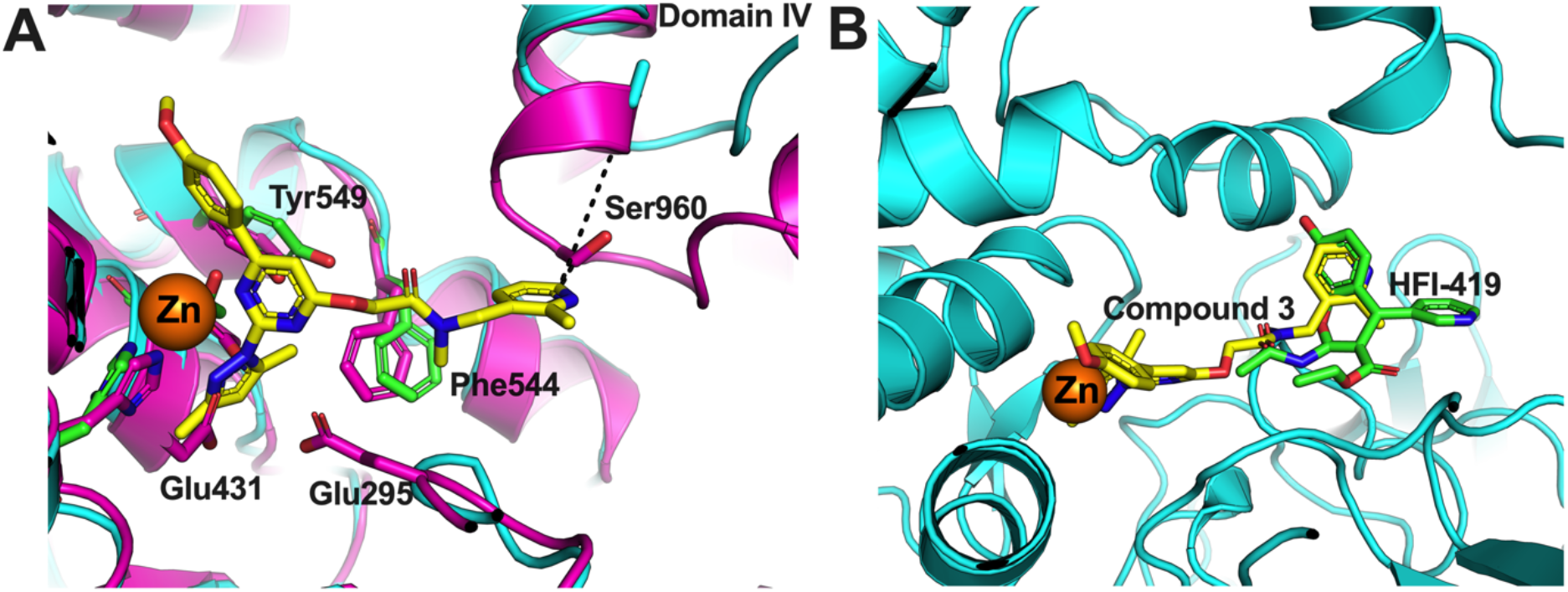
**Panel A**, comparison of the structure of IRAP (cyan cartoon) with compound **3** (yellow sticks) with the closed conformation of IRAP crystallized with a transition-state analogue (magenta cartoon, PDB ID 5MJ6). Dotted line indicates the distance between the 2-methylpyridine group of compound **3** with Ser960 of domain IV of IRAP in the open (7.6Å) and closed (1.4Å) conformations. This compound configuration would be incompatible with the closed conformation of IRAP due to steric clashes with domain IV. Residues in the catalytic site that differ in orientation between the two IRAP conformations are shown as sticks (green sticks for the open conformation and magenta sticks for the closed conformation). **Panel B**, superposition of the IRAP/compound **3** structure with the IRAP/HFI-419 structure indicates that HFI-419 binds at a location where the 2-methylpyridine group of compound **3** also binds.

**Figure 7:**
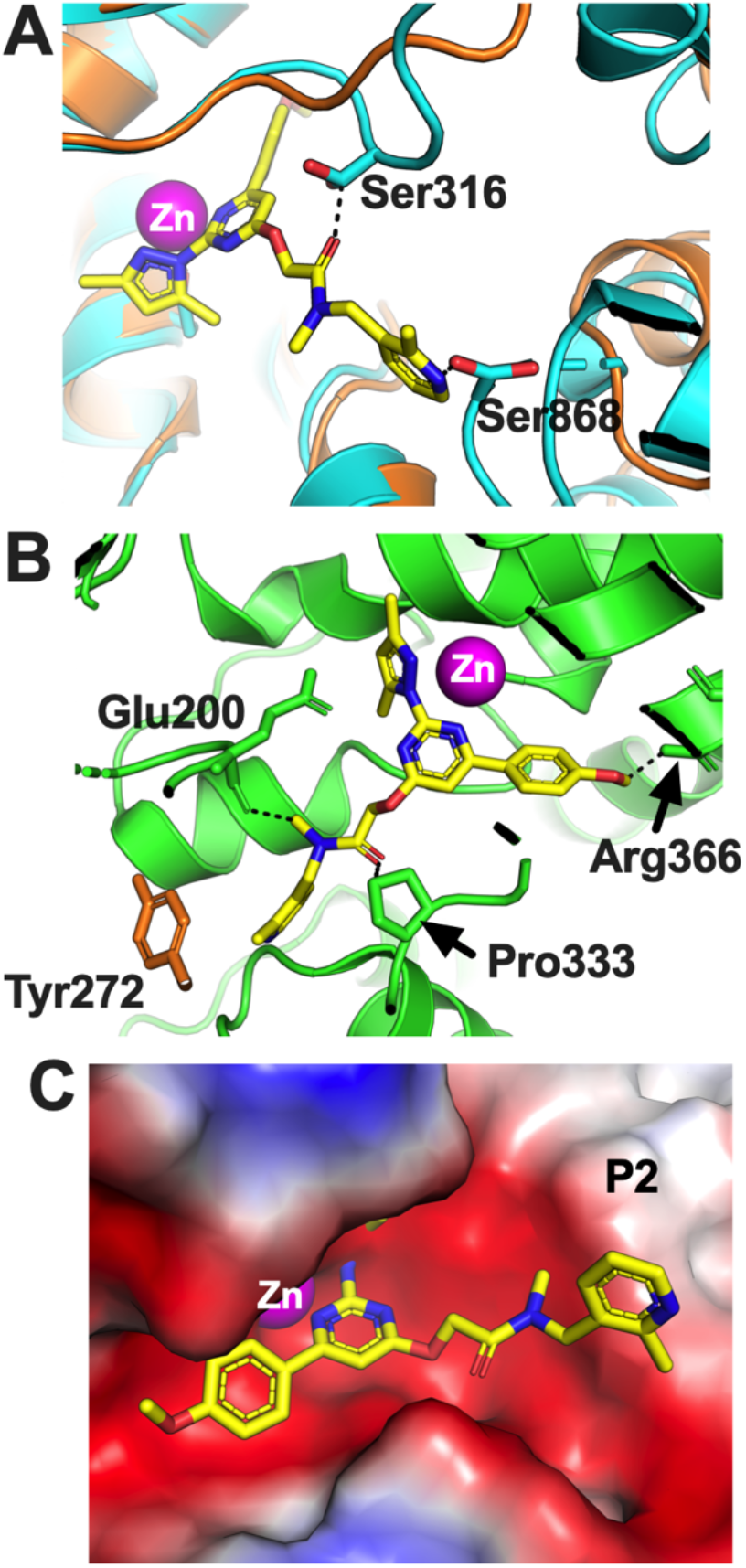
Structural basis for observed selectivity of compound **3. Panel A**, superimposing the crystal structure of the closed conformation of ERAP1 (PDB ID 6Q4R, in cyan) to the IRAP-compound **3** structure (IRAP as an orange cartoon, compound **3** shown in yellow sticks) reveals potential steric clashes with Ser316 of the ERAP1 GAMEN loop and Ser868 of the ERAP1 domain IV. **Panel B**, superimposing the crystal structure of the only known conformation of ERAP2^34^ (PDB ID 5AB0, in green) with the structure of IRAP/compound **3** (yellow sticks) reveals putative clashes between the inhibitor and ERAP2 with Pro333 of the GAMEN loop, Arg366 and Glu200. **Panel C**, superimposing the crystal structure of the open conformation of ERAP1 (PDB ID 3MDJ, shown in surface representation colored by electrostatic potential) to the IRAP-compound **3** structure (compound **3** shown in yellow sticks) reveals the lack of formation of the P2 pocket (compare to Figure 4D) that accommodates the 2-methylpyridine group of compound **3**.

Tyr549 is a key active site residue that promotes catalysis by stabilizing the tetrahedral intermediate formed after nucleophilic attack of a water molecule to the carbonyl of the scissile peptide bond^32^. The relative configuration of the Tyr549 sidechain has been observed to be different in the empty versus ligand-bound crystal structures of IRAP and this has been hypothesized to relate to the higher catalytic efficiency of the closed state^19^. Indeed, Tyr549 is oriented away from the catalytic Zn(II) atom in the IRAP/compound **3** structure, suggesting that compound **3** binding helps retain IRAP in a catalytically inactive state.

Compound **3** was found to have exceptional selectivity versus the other two members of the oxytocinase sub-family of M1 aminopeptidases, ERAP1 and ERAP2 (Figure 1A). To understand this selectivity, we aligned the IRAP/compound **3** structure with ERAP1 and ERAP2 structures based on their domain I (Figure 7). We utilized both experimentally determined conformations of ERAP1^33^. Visual inspection of the aligned structures revealed atomic interactions that can readily explain the observed selectivity. The observed orientation of compound **3** in IRAP, would be incompatible with the known closed conformation of ERAP1 due to clashes with the non-conserved Ser316 of the ERAP1 GAMEN loop and Ser868 of the ERAP1 domain IV (Figure 7A). Similarly, the non-conserved Pro333 of the ERAP2 GAMEN loop, as well as side chains of Glu200 and Arg366 are likely sterically non-compatible with the observed compound **3** configuration (Figure 7B). Finally, although no steric clashes can be predicted between compound **3** and the open conformation of ERAP1, lack of a P2 pocket would be expected to weaken binding (Figure 7C). Thus, a series of enzyme-specific atomic clashes or lack of interactions appears sufficient to explain the exceptional selectivity of compound **3** for IRAP.

## DISCUSSION

Here, we describe the characterization and analysis of the mechanism of action of a novel zinc-aminopeptidase inhibitor that utilizes a pyrazolylpyrimidine scaffold to chelate the active site zinc(II). While many zinc(II) chelating groups have been utilized as pharmacophores for metallopeptidase inhibition^35^, including the structurally similar pyridinyl pyrimidine scaffold^36^, the particular combination of the 2-(pyrazol-1-yl)pyrimidine appears to be structurally suited for the active site of IRAP. While this compound appears to have high selectivity for IRAP due to its interplay with structural features in the open conformation of the enzyme, the pyrazolylpyrimidine scaffold may also be useful in designing inhibitors for other members of the M1 family of aminopeptidases.

Although the uncompetitive mechanism of action discovered for compound **3** was, at first examination, unexpected, such a mechanism is not the first time reported for IRAP. Indeed, our group recently reported an uncompetitive mechanism for another compound, namely the widely used and commercially available IRAP inhibitor named HFI-419^21^. This compound however was found to bind to an allosteric site, formed only in the open conformation of IRAP, and an uncompetitive mechanism of action is not uncommon for allosteric inhibitors. The uniqueness of compound **3** however is that it binds to the active site and still displays uncompetitive/noncompetitive kinetics. This behaviour has been observed before for enzymes that cycle through multiple states, and inhibitor binding to only a single conformation can result in apparent non-competitive kinetics^37,38^. Thus conformational cycling between two or more conformational states, as proposed for IRAP, can help explain this behaviour^19^. Indeed, enzymatic mechanisms that involve conformational cycling have been proposed for homologous M1 aminopeptidases such as ERAP1^33,39^, aminopeptidase N^40,41^ and lanthipeptide zinc-metallopeptidase^42^.

While stabilization of an open, less-active, conformation of IRAP is an appealing mechanism of action, it by itself appears insufficient to fully explain all our experimental observations, and specifically the very low potency of compound **3** for blocking the cleavage of large peptidic substrates including oxytocin. Although a similar pattern of inhibitor activity was observed for the allosteric inhibitor HFI-419, in that case the potential of concurrent occupancy of the active and allosteric sites allowed the formulation of a simple kinetic model that explained inhibitor activity, based on the hypothesis that the open conformation is sufficiently active to hydrolyze oxytocin^21^. Here, in contrast, the binding of compound **3** in the active site, precludes this explanation and therefore brings into doubt its generality, necessitating an alternate model. One possible solution to this paradox may rely on the large conformational lability of the subfamily of oxytocinases of M1 aminopeptidases as revealed by X-ray crystallography and molecular dynamics^43^. Indeed, the highly homologous ERAP1, an aminopeptidase specialized for long peptides^33^, has been crystallized in a significantly more “open” conformation that is suitable for binding large substrates. Such a conformation (termed “fully open”) is theoretically accessible for IRAP, based on molecular dynamics calculations^43^. In this context, a possible model of IRAP substrate trimming and inhibition is shown in Figure 8. According to this model, small substrates induce the transition from the open conformation E to the closed E^*^, which promotes their catalytic turnover. Due to lack of direct access to external solvent, after catalysis, the enzyme must cycle to the open conformation E to allow product release and new substrate capture. Larger substrates need a “fully open” conformation to bind, E ^#^, and repeating catalytic cycles can proceed in that state. Compound **3** cannot bind to the closed conformation due to steric clashes but binds to the open conformation with high affinity (by design) and possibly to the fully open conformation with lower affinity, due to the lack of a well-formed P2 pocket (Figure 7). Similarly, allosteric inhibitors such as HFI-419 can only bind to the open conformation and are thus ineffective versus large substrates^21^. Medium-sized substrates, like the 9mer peptide, may lie in the middle region and utilize both the open and fully open conformation resulting in intermediate or biphasic efficiency. Since the fully open conformation has not been observed yet experimentally, further structural work focusing on the larger substrates and specifically on the cyclic peptide substrates will be necessary to provide further support for this hypothesis.

**Figure 8:**
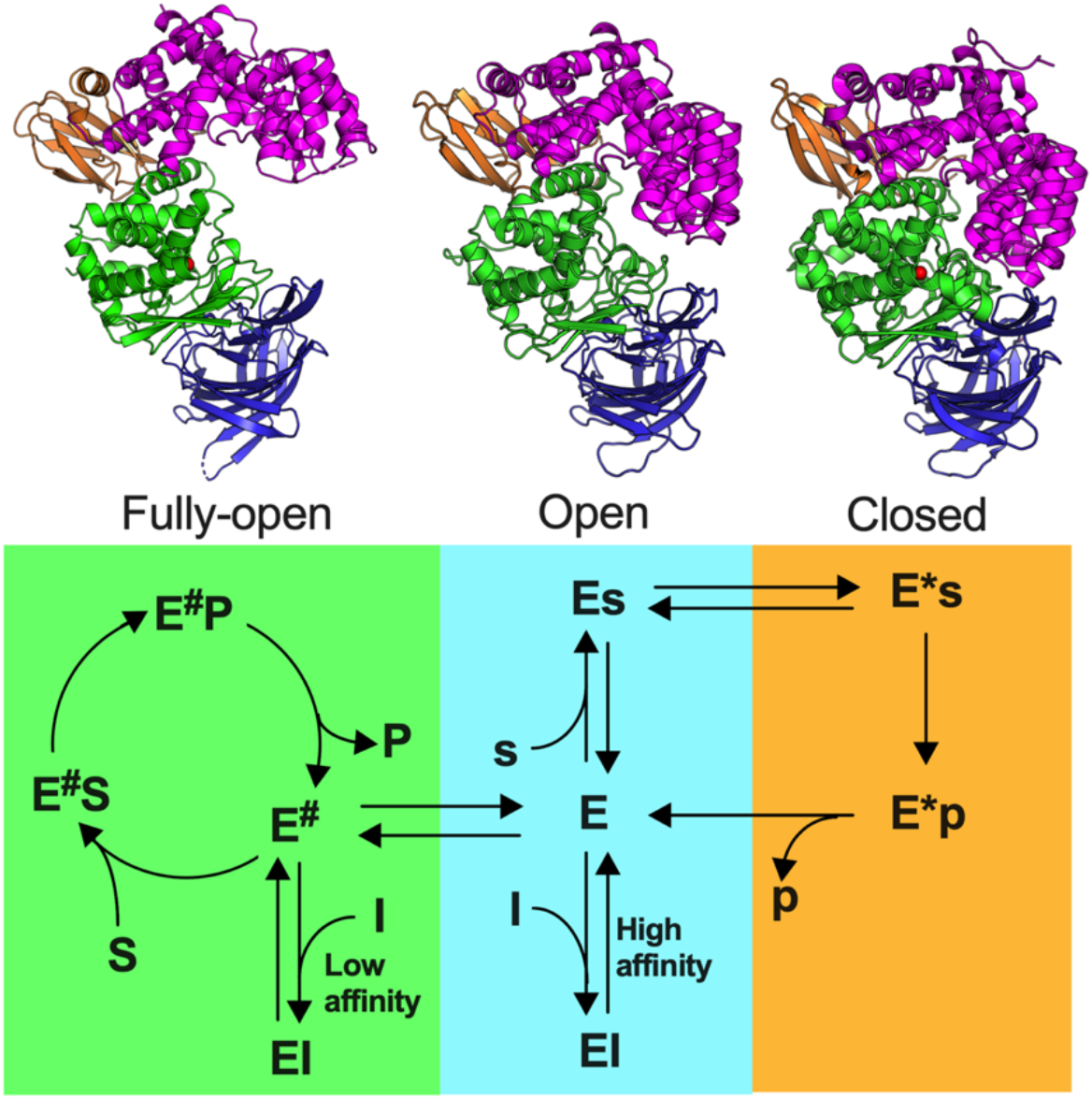
Schematic representation of putative IRAP catalytic cycles. **E, E**^**\***^ and **E** ^**#**^ represent different conformations of the enzyme: **E** is the experimentally observed open conformation (PDB ID: 8P0I), **E**^**\***^ is the experimentally observed closed conformation (PDB ID: 5MJ6) and **E**^**#**^ is a speculative “fully” open conformation, resembling the ERAP1 open conformation (PDB ID: 3MDJ). Small substrates (**s**) are processed after binding to the **E** conformation and inducing the transition to the closed **E**^**\***^ but need to cycle back to the **E** conformation to allow for product release and new substrate capture. Larger substrates (**S**) can only be bound by the fully open conformation **E**^**#**^ which can perform catalytic turnover possibly without needing to access closed states. **I** indicates an inhibitor (i.e., compound **3**) that recognizes the **E** conformation with high-affinity and the E ^#^ conformation with low affinity.

The development of enzyme inhibitors as drug candidates is a lengthy and arduous process that relies, from early on, on the availability of physiologically relevant yet easily implementable *in vitro* assays both for high-throughput screening and hit-to-lead optimization. Small chromogenic or fluorogenic dipeptide-like substrates are widely used for drug discovery purposes in the field of peptidases and for aminopeptidases. Here, we successfully optimized an IRAP inhibitor both for potency and selectivity guided by the widely used Leu-AMC assay. This optimization, however, resulted in a complex potency landscape for physiologically relevant larger peptides, exposing gaps in knowledge on how those substrates are processed. Although this may be due to the intricacies of the IRAP catalytic cycle, it constitutes a cautionary tale in favor of using more physiologically relevant orthogonal assays early in the drug discovery process. Indeed, a high-throughput screen for the discovery of inhibitors of the highly homologous aminopeptidase ERAP1, resulted in the discovery of compounds that activate Leu-AMC cleavage but inhibit larger peptide trimming^44^. Thus, the application of appropriate orthogonal assays is critical for both drug discovery and for understanding the mechanism of function of enzymes that operate with large conformational rearrangements.

Our findings have important repercussions on the preclinical development of inhibitors of IRAP. The low efficacy of compound **3** in blocking the processing of large or cyclic peptides may limit its value for further pre-clinical development that focuses on the biological functions of IRAP that relate to the degradation of such substrates. Still, compound **3** is effective in blocking the trimming of small substrates, with potent, albeit partial, activity for a 9mer antigenic peptide. IRAP is highly active in trimming peptides of 9 amino acids or smaller, in sharp contrast to its homologous ERAP1^18^. When trimming smaller peptides, IRAP may play physiological roles in processing antigenic epitopes in Dendritic Cells during the process of cross-presentation^4,45^. It may also contribute to the generation of smaller-size epitopes such as the SL8 in the case of ovalbumin cross-presentation by mouse cells, as indicated by our findings. This capacity may be less favourable to antigen cross-presentation by human MHC-I molecules that generally prefer 9-mer peptides. In this context, compound **3**, may hold value for developing inhibitors of IRAP that regulate antigen presentation by selectively limiting antigenic epitope destruction while allowing elongated precursor processing. Substrate-selective inhibition in aminopeptidases has been observed before in systems with clinical relevance: we recently described an allosteric regulator of ERAP1 that shapes the immunopeptidome of cells (the repertoire of presented peptides) in unique patterns^46^. Additional work is necessary to explore the potential value of this compound in regulating antigen presentation and the immunopeptidome of cells.

In summary, we describe a novel, highly selective, small-molecule IRAP inhibitor that uses an unconventional mechanism of action by binding to the active site of the open conformation of the enzyme. Our results emphasize that conformational changes, both in the overall configuration of the protein domains and configuration of structural elements of the active site, are central to the IRAP catalytic cycle and that conformational restriction may constitute a novel approach to modulating this enzyme’s activity. It also shows limitations in our understanding of how IRAP recognizes and processes some of its substrates. Additional structural analyses of IRAP with cyclic or other large peptide substrates or analogues will be necessary to address these questions and to help guide the design of new compounds with the exciting prospect of selectively targeting IRAP’s biological functions, including the possibility of substrate-specific modulation of the enzyme’s activity. Any optimization of such compounds for potential clinical use will require an understanding of the physiological substrates that drive efficacy and incorporation of native substrate cleavage inhibition assays within the project’s toolbox.

## Author Contributions

A.M. produced and purified recombinant enzymes, performed enzymatic analysis, analyzed and interpreted data, crystallized the protein, and solved the crystal structure. A.B., A.D.F., J.S.F. and D.W.H. conceived, designed, and supervised the small molecule discovery and optimization project. G.G. performed enzymatic analysis, analyzed and interpreted data. G.G., F.X.M. and P.vE. designed and performed the cross-presentation assay. P.G. helped with protein crystallization, performed X-ray diffraction experiments and analyzed X-ray diffraction data. E.S. conceived and supervised the structural biology and enzymology project, analyzed and interpreted data, and wrote the manuscript with input from all authors. All authors have approved the final version of the manuscript.

## Funding

Funding was provided by internal funds of the National Centre for Scientific Research “Demokritos” and by the project “The Greek Research Infrastructure for Personalized Medicine (pMedGR)” (MIS 5002802), which is implemented under the Action “Reinforcement of the Research and Innovation Infrastructure,” funded by the Operational Programme “Competitiveness, Entrepreneurship and Innovation” (NSRF 2014–2020) and co-financed by Greece and the European Union (European Regional Development Fund). Funding was also provided by the European Commission in the context of the Marie Skłodowska-Curie Action European Training Network CAPSTONE (954992 – CAPSTONE – H2020-MSCA-ITN-2020). The authors would also like to acknowledge financial support from Pharmaxis Ltd (now Syntara Limited).

## Acknowledgments

We thank the beamline scientists of EMBL-Hamburg for their assistance during data collection at the P13 beamline of Petra III, Hamburg, Germany and Angelique Greco and Graeme Stevenson for their help with the virtual screen.

## Conflict of interest

The authors declare no competing financial interest. A.B., A.D.F., J.S.F. and D.W.H. were employees and shareholders of Pharmaxis Ltd at the time this work was carried out.

## Data deposition

Atomic coordinates and structure factors have been deposited in the Protein Data Bank (https://www.rcsb.org/) with PDB code 8P0I.

## Notes

### Competing Interest Statement

The authors have declared no competing interest.

## References

(1) Tsujimoto, M.; Hattori, A. The Oxytocinase Subfamily of M1 Aminopeptidases. Biochim Biophys Acta 2005, 1751 (1), 9–18.

(2) Weimershaus, M.; Mauvais, F.-X.; Evnouchidou, I.; Lawand, M.; Saveanu, L.; van Endert, P. IRAP Endosomes Control Phagosomal Maturation in Dendritic Cells. Front. Cell Dev. Biol. 2020, 8, 1419. 10.3389/fcell.2020.585713.

(3) Jordens, I.; Molle, D.; Xiong, W.; Keller, S. R.; McGraw, T. E. Insulin-Regulated Aminopeptidase Is a Key Regulator of GLUT4 Trafficking by Controlling the Sorting of GLUT4 from Endosomes to Specialized Insulin-Regulated Vesicles. Mol. Biol. Cell 2010, 21 (12), 2034–2044. 10.1091/mbc.e10-02-0158.

(4) Saveanu, L. IRAP Identifies an Endosomal Compartment Required for MHC Class I Cross-Presentation. Science 2009, 325, 213–217. 10.1126/science.1172845.

(5) Tsujimoto, M.; Mizutani, S.; Adachi, H.; Kimura, M.; Nakazato, H.; Tomoda, Y. Identification of Human Placental Leucine Aminopeptidase as Oxytocinase. Arch. Biochem. Biophys. 1992, 292 (2), 388–392. 10.1016/0003-9861(92)90007-J.

(6) Herbst, J. J.; Ross, S. A.; Scott, H. M.; Bobin, S. A.; Morris, N. J.; Lienhard, G. E.; Keller, S. R. Insulin Stimulates Cell Surface Aminopeptidase Activity toward Vasopressin in Adipocytes. Am J Physiol 1997, 272 (4 Pt 1), E600–6. 10.1152/ajpendo.1997.272.4.E600.

(7) Albiston, A. L.; Morton, C. J.; Ng, H. L.; Pham, V. V.; Yeatman, H. R.; Ye, S.; Fernando, R. N.; De Bundel, D.; Ascher, D. B.; Mendelsohn, F. A. O.; Parker, M. W.; Chai, S. Y. Identification and Characterization of a New Cognitive Enhancer Based on Inhibition of Insulin-Regulated Aminopeptidase. FASEB J. 2008, 22 (12), 4209–4217. 10.1096/fj.08-112227.

(8) Chai, S. Y.; Yeatman, H. R.; Parker, M. W.; Ascher, D. B.; Thompson, P. E.; Mulvey, H. T.; Albiston, A. L. Development of Cognitive Enhancers Based on Inhibition of Insulin-Regulated Aminopeptidase. BMC Neurosci. 2008, 9 Suppl 2, S14. 10.1186/1471-2202-9-S2-S14.

(9) Gaspari, T.; Matthew Shen; Wang, Y.; Shastry, A.; Chai, S. Y.; Samuel, C.; Widdop, R. A9684 Comparing Anti-Fibrotic Effects of the Irap Inhibitor, Hfi-419 to an Angiotensin Receptor Blocker and Ace Inhibitor in a High Salt-Induced Mouse Model of Kidney Disease. J. Hypertens. 2018, 36, e56. 10.1097/01.hjh.0000548218.38141.2a.

(10) Gaspari, T.; Lee, H. W.; Fan, K.; Salimova, E.; Spizzo, I.; Samuel, C.; Abouelkheir, M.; Thompson, P.; Chai, S. Y.; Widdop, R. A9871 Insulin Regulated Aminopeptidase (Irap) Inhibition Completely Reverses Age-Induced Cardiac Fibrosis and Improves Cardiac Function. J. Hypertens. 2018, 36, e57. 10.1097/01.hjh.0000548219.38141.63.

(11) Georgiadis, D.; Ziotopoulou, A.; Kaloumenou, E.; Lelis, A.; Papasava, A. The Discovery of Insulin-Regulated Aminopeptidase (IRAP) Inhibitors: A Literature Review. Front. Pharmacol. 2020, 11, 1433. 10.3389/fphar.2020.585838.

(12) Gising, J.; Honarnejad, S.; Bras, M.; Baillie, G. L.; McElroy, S. P.; Jones, P. S.; Morrison, A.; Beveridge, J.; Hallberg, M.; Larhed, M. The Discovery of New Inhibitors of Insulin-Regulated Aminopeptidase by a High-Throughput Screening of 400,000 Drug-like Compounds. Int. J. Mol. Sci. 2024, 25 (7), 4084. 10.3390/ijms25074084.

(13) Engen, K.; Lundbäck, T.; Yadav, A.; Puthiyaparambath, S.; Rosenström, U.; Gising, J.; Jenmalm-Jensen, A.; Hallberg, M.; Larhed, M. Inhibition of Insulin-Regulated Aminopeptidase by Imidazo [1,5-α]Pyridines-Synthesis and Evaluation. Int. J. Mol. Sci. 2024, 25 (5), 2516. 10.3390/ijms25052516.

(14) Seyer, B.; Diwakarla, S.; Burns, P.; Hallberg, A.; Grönbladh, A.; Hallberg, M.; Chai, S. Y. Insulin-Regulated Aminopeptidase Inhibitor-Mediated Increases in Dendritic Spine Density Are Facilitated by Glucose Uptake. J. Neurochem. 2020, 153 (4), 485–494. 10.1111/JNC.14880.

(15) El-Hawli, A.; Qaradakhi, T.; Hayes, A.; Rybalka, E.; Smith, R.; Caprnda, M.; Opatrilova, R.; Gazdikova, K.; Benckova, M.; Kruzliak, P.; Zulli, A. IRAP Inhibition Using HFI419 Prevents Moderate to Severe Acetylcholine Mediated Vasoconstriction in a Rabbit Model. Biomed Pharmacother 2017, 86, 23–26. 10.1016/j.biopha.2016.11.142.

(16) Krskova, K.; Balazova, L.; Dobrocsyova, V.; Olszanecki, R.; Suski, M.; Chai, S. Y.; Zorad, Š. Insulin-Regulated Aminopeptidase Inhibition Ameliorates Metabolism in Obese Zucker Rats. Front. Mol. Biosci. 2020, 7, 383. 10.3389/fmolb.2020.586225.

(17) Hermans, S. J.; Ascher, D. B.; Hancock, N. C.; Holien, J. K.; Michell, B. J.; Chai, S. Y.; Morton, C. J.; Parker, M. W. Crystal Structure of Human Insulin-Regulated Aminopeptidase with Specificity for Cyclic Peptides. Protein Sci 2015, 24 (2), 190–199. 10.1002/pro.2604.

(18) Mpakali, A.; Saridakis, E.; Harlos, K.; Zhao, Y.; Papakyriakou, A.; Kokkala, P.; Georgiadis, D.; Stratikos, E. Crystal Structure of Insulin-Regulated Aminopeptidase with Bound Substrate Analogue Provides Insight on Antigenic Epitope Precursor Recognition and Processing. J. Immunol. Baltim. Md 1950 2015, 195 (6), 2842–2851. 10.4049/jimmunol.1501103.

(19) Mpakali, A.; Saridakis, E.; Harlos, K.; Zhao, Y.; Kokkala, P.; Georgiadis, D.; Giastas, P.; Papakyriakou, A.; Stratikos, E. Ligand-Induced Conformational Change of Insulin-Regulated Aminopeptidase: Insights on Catalytic Mechanism and Active Site Plasticity. J. Med. Chem. 2017, 60 (7), 2963–2972. 10.1021/acs.jmedchem.6b01890.

(20) Mpakali, A.; Saridakis, E.; Giastas, P.; Maben, Z.; Stern, L. J.; Larhed, M.; Hallberg, M.; Stratikos, E. Structural Basis of Inhibition of Insulin-Regulated Aminopeptidase by a Macrocyclic Peptidic Inhibitor. ACS Med. Chem. Lett. 2020, 11 (7), 1429–1434. 10.1021/acsmedchemlett.0c00172.

(21) Mpakali, A.; Barla, I.; Lu, L.; Ramesh, K. M.; Thomaidis, N.; Stern, L. J.; Giastas, P.; Stratikos, E. Mechanisms of Allosteric Inhibition of Insulin-Regulated Aminopeptidase. J. Mol. Biol. 2024, 436 (6), 168449. 10.1016/j.jmb.2024.168449.

(22) Stamogiannos, A.; Maben, Z.; Papakyriakou, A.; Mpakali, A.; Kokkala, P.; Georgiadis, D.; Stern, L. J.; Stratikos, E. Critical Role of Interdomain Interactions in the Conformational Change and Catalytic Mechanism of Endoplasmic Reticulum Aminopeptidase 1. Biochemistry 2017, 56 (10), 1546–1558. 10.1021/acs.biochem.6b01170.

(23) Temponeras, I.; Chiniadis, L.; Papakyriakou, A.; Stratikos, E. Discovery of Selective Inhibitor Leads by Targeting an Allosteric Site in Insulin-Regulated Aminopeptidase. Pharm. Basel Switz. 2021, 14 (6), 584. 10.3390/ph14060584.

(24) Atagunduz, P.; Appel, H.; Kuon, W.; Wu, P.; Thiel, A.; Kloetzel, P.-M.; Sieper, J. HLA-B27-Restricted CD8+ T Cell Response to Cartilage-Derived Self Peptides in Ankylosing Spondylitis. Arthritis Rheum. 2005, 52 (3), 892–901. 10.1002/art.20948.

(25) Shankar, P.; Russo, M.; Harnisch, B.; Patterson, M.; Skolnik, P.; Lieberman, J. Impaired Function of Circulating HIV-Specific CD8(+) T Cells in Chronic Human Immunodeficiency Virus Infection. Blood 2000, 96 (9), 3094–3101.

(26) Steiner, Q.-G.; Otten, L. A.; Hicks, M. J.; Kaya, G.; Grosjean, F.; Saeuberli, E.; Lavanchy, C.; Beermann, F.; McClain, K. L.; Acha-Orbea, H. In Vivo Transformation of Mouse Conventional CD8alpha+ Dendritic Cells Leads to Progressive Multisystem Histiocytosis. Blood 2008, 111 (4), 2073–2082. 10.1182/blood-2007-06-097576.

(27) Karttunen, J.; Sanderson, S.; Shastri, N. Detection of Rare Antigen-Presenting Cells by the lacZ T-Cell Activation Assay Suggests an Expression Cloning Strategy for T-Cell Antigens. Proc. Natl. Acad. Sci. U. S. A. 1992, 89 (13), 6020–6024. 10.1073/pnas.89.13.6020.

(28) Albiston, A. L.; Morton, C. J.; Ng, H. L.; Pham, V.; Yeatman, H. R.; Ye, S.; Fernando, R. N.; De Bundel, D.; Ascher, D. B.; Mendelsohn, F. A.; Parker, M. W.; Chai, S. Y. Identification and Characterization of a New Cognitive Enhancer Based on Inhibition of Insulin-Regulated Aminopeptidase. Faseb J 2008, 22 (12), 4209–4217. https://doi.org/fj.08-112227 [pii] 10.1096/fj.08-112227.

(29) Firat, E.; Saveanu, L.; Aichele, P.; Staeheli, P.; Huai, J.; Gaedicke, S.; Nil, A.; Besin, G.; Kanzler, B.; van Endert, P.; Niedermann, G. The Role of Endoplasmic Reticulum-Associated Aminopeptidase 1 in Immunity to Infection and in Cross-Presentation. J Immunol 2007, 178 (4), 2241–2248.

(30) Albiston, A. L.; Pham, V.; Ye, S.; Ng, L.; Lew, R. A.; Thompson, P. E.; Holien, J. K.; Morton, C. J.; Parker, M. W.; Chai, S. Y. Phenylalanine-544 Plays a Key Role in Substrate and Inhibitor Binding by Providing a Hydrophobic Packing Point at the Active Site of Insulin-Regulated Aminopeptidase. Mol. Pharmacol. 2010, 78 (4), 600–607. 10.1124/mol.110.065458.

(31) Vourloumis, D.; Mavridis, I.; Athanasoulis, A.; Temponeras, I.; Koumantou, D.; Giastas, P.; Mpakali, A.; Magrioti, V.; Leib, J.; van Endert, P.; Stratikos, E.; Papakyriakou, A. Discovery of Selective Nanomolar Inhibitors for Insulin-Regulated Aminopeptidase Based on α-Hydroxy-β-Amino Acid Derivatives of Bestatin. J. Med. Chem. 2022, 65 (14), 10098–10117. 10.1021/acs.jmedchem.2c00904.

(32) Tholander, F.; Muroya, A.; Roques, B. P.; Fournié-Zaluski, M. C.; Thunnissen, M. M. G. M.; Haeggström, J. Z. Structure-Based Dissection of the Active Site Chemistry of Leukotriene A4 Hydrolase: Implications for M1 Aminopeptidases and Inhibitor Design. Chem. Biol. 2008, 15 (9), 920–929. 10.1016/j.chembiol.2008.07.018.

(33) Kochan, G.; Krojer, T.; Harvey, D.; Fischer, R.; Chen, L.; Vollmar, M.; von Delft, F.; Kavanagh, K. L.; Brown, M. A.; Bowness, P.; Wordsworth, P.; Kessler, B. M.; Oppermann, U. Crystal Structures of the Endoplasmic Reticulum Aminopeptidase-1 (ERAP1) Reveal the Molecular Basis for N-Terminal Peptide Trimming. Proc. Natl. Acad. Sci. U. S. A. 2011, 108 (19), 7745–7750. 10.1073/pnas.1101262108.

(34) Mpakali, A.; Giastas, P.; Mathioudakis, N.; Mavridis, I. M.; Saridakis, E.; Stratikos, E. Structural Basis for Antigenic Peptide Recognition and Processing by Endoplasmic Reticulum (ER) Aminopeptidase 2. J. Biol. Chem. 2015, 290 (43), 26021–26032. 10.1074/jbc.M115.685909.

(35) Chen, A. Y.; Adamek, R. N.; Dick, B. L.; Credille, C. V.; Morrison, C. N.; Cohen, S. M. Targeting Metalloenzymes for Therapeutic Intervention. Chem. Rev. 2019, 119 (2), 1323–1455. 10.1021/acs.chemrev.8b00201.

(36) Hu, X.; Addlagatta, A.; Matthews, B. W.; Liu, J. O. Identification of Pyridinylpyrimidines as Inhibitors of Human Methionine Aminopeptidases. Angew. Chem. Int. Ed Engl. 2006, 45 (23), 3772–3775. 10.1002/anie.200600757.

(37) Blat, Y. Non-Competitive Inhibition by Active Site Binders. Chem. Biol. Drug Des. 2010, 75 (6), 535–540. 10.1111/j.1747-0285.2010.00972.x.

(38) Pesaresi, A. Mixed and Non-Competitive Enzyme Inhibition: Underlying Mechanisms and Mechanistic Irrelevance of the Formal Two-Site Model. J. Enzyme Inhib. Med. Chem. 2023, 38 (1), 2245168. 10.1080/14756366.2023.2245168.

(39) Maben, Z.; Arya, R.; Georgiadis, D.; Stratikos, E.; Stern, L. J. Conformational Dynamics Linked to Domain Closure and Substrate Binding Explain the ERAP1 Allosteric Regulation Mechanism. Nat. Commun. 2021, 12 (1), 5302. 10.1038/s41467-021-25564-w.

(40) Addlagatta, A.; Gay, L.; Matthews, B. W. Structural Basis for the Unusual Specificity of Escherichia Coli Aminopeptidase N. Biochemistry 2008, 47 (19), 5303–5311. 10.1021/bi7022333.

(41) Chen, L.; Lin, Y. L.; Peng, G.; Li, F. Structural Basis for Multifunctional Roles of Mammalian Aminopeptidase N. Proc Natl Acad Sci U A 2012, 109 (44), 17966–17971. 10.1073/pnas.12101231091210123109 [pii].

(42) Zhao, C.; Sheng, W.; Wang, Y.; Zheng, J.; Xie, X.; Liang, Y.; Wei, W.; Bao, R.; Wang, H. Conformational Remodeling Enhances Activity of Lanthipeptide Zinc-Metallopeptidases. Nat. Chem. Biol. 2022, 18 (7), 724–732. 10.1038/s41589-022-01018-2.

(43) Papakyriakou, A.; Stratikos, E. The Role of Conformational Dynamics in Antigen Trimming by Intracellular Aminopeptidases. Front. Immunol. 2017, 8, 946. 10.3389/fimmu.2017.00946.

(44) Liddle, J.; Hutchinson, J. P.; Kitchen, S.; Rowland, P.; Neu, M.; Cecconie, T.; Holmes, D. S.; Jones, E.; Korczynska, J.; Koumantou, D.; Lea, J. D.; Nickels, L.; Pemberton, M.; Phillipou, A.; Schneck, J. L.; Sheehan, H.; Tinworth, C. P.; Uings, I.; Wojno-Picon, J.; Young, R. J.; Stratikos, E. Targeting the Regulatory Site of ER Aminopeptidase 1 Leads to the Discovery of a Natural Product Modulator of Antigen Presentation. J. Med. Chem. 2020, 63 (6), 3348–3358. 10.1021/acs.jmedchem.9b02123.

(45) Georgiadou, D.; Hearn, A.; Evnouchidou, I.; Chroni, A.; Leondiadis, L.; York, I. A.; Rock, K. L.; Stratikos, E. Placental Leucine Aminopeptidase Efficiently Generates Mature Antigenic Peptides in Vitro but in Patterns Distinct from Endoplasmic Reticulum Aminopeptidase 1. J. Immunol. Baltim. Md 1950 2010, 185 (3), 1584–1592. 10.4049/jimmunol.0902502.

(46) Temponeras, I.; Samiotaki, M.; Koumantou, D.; Nikopaschou, M.; Kuiper, J. J. W.; Panayotou, G.; Stratikos, E. Distinct Modulation of Cellular Immunopeptidome by the Allosteric Regulatory Site of ER Aminopeptidase 1. Eur. J. Immunol. 2023, e2350449. 10.1002/eji.202350449.

