## Supporting Figure 1 for "Stabilization of the open conformation of Insulin-Regulated Aminopeptidase by a novel substrate-selective small molecule inhibitor"

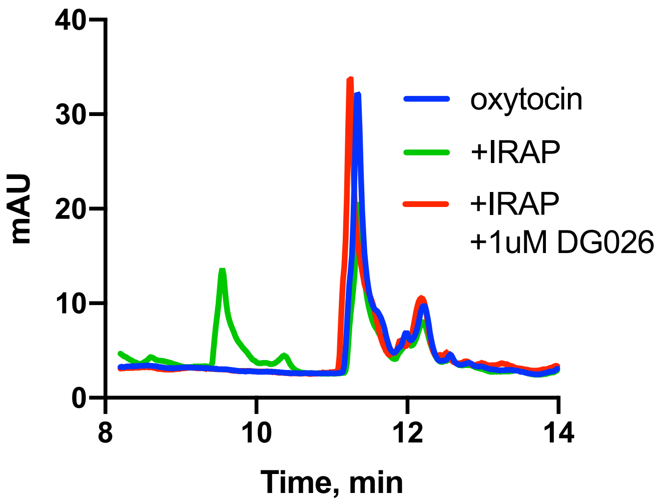


**Supporting Figure 1:** Reversed-phase HPLC chromatograms of oxytocin incubated with 10 nM IRAP for 30 min at 37°C in the presence or absence of the transition-state analogue inhibitor DG026.

**Preparation of Compound 2**

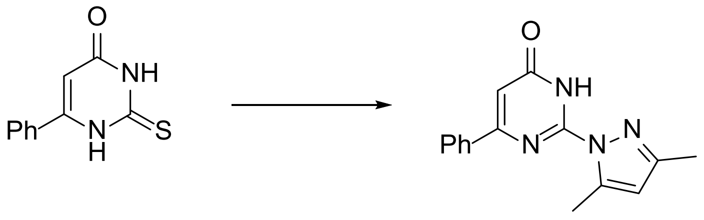


**a) Preparation of 2-(3,5-dimethylpyrazol-1-yl)-4-phenyl-1*H*-pyrimidin-6-one**

A mixture of 6-phenyl-2-thioxo-1*H*-pyrimidin-4-one (0.41 g, 2 mmol) and hydrazine monohydrate (1.5 mL, 30 mmol) was heated at 90 °C for 2 hours. A crystalline solid crashed out. The reaction mixture was cooled to room temperature and diluted with water (10 mL). The product was filtered, washed with water. The solid was transferred to a round bottom flask and dissolved in a mixture of ethanol (2 mL) and acetic acid (3 mL). Pentane-2,4-dione (0.21 mL, 2 mmol) was added and the mixture was heated at reflux for two hours. The reaction mixture was concentrated under vacuum and diluted with water (10 mL). The product was collected by filtration and dried in an oven at 60 °C when the title compound was obtained as a white solid (0.43 g, 1.6 mmol, 81%). $^1$H NMR (300 MHz, Chloroform-*d*) δ 10.41 (s, 1H), 8.04 – 7.96 (m, 2H), 7.56 – 7.47 (m, 3H), 6.71 (s, 1H), 6.11 (d, *J* = 1.1 Hz, 1H), 2.84 (d, *J* = 0.9 Hz, 3H), 2.30 (s, 3H).

**b) Preparation of *tert*-butyl 2-[2-(3,5-dimethylpyrazol-1-yl)-6-phenyl-pyrimidin-4-yl]oxyacetate**

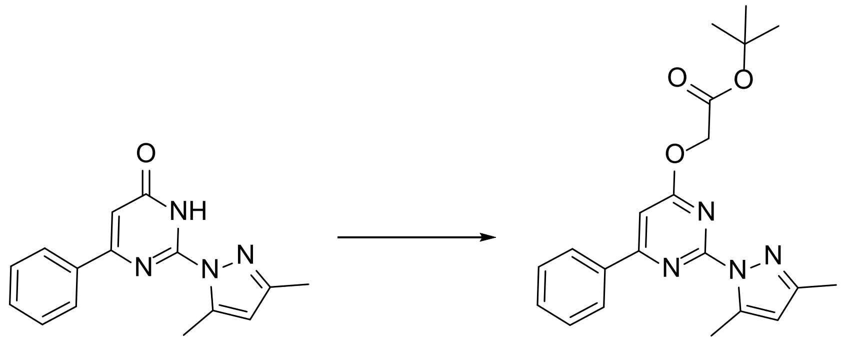


*tert*-Butyl bromoacetate (1.3 mL, 8.9 mmol) was added to a stirring mixture of 2-(3,5-dimethylpyrazol-1-yl)-4-phenyl-1*H*-pyrimidin-6-one (1.8 g, 6.9 mmol) and caesium carbonate (2.9 g, 8.9 mmol) in DMF (10 mL). The mixture was stirred at room temperature for two hours. Water (50 mL) was added and the product was extracted with ethyl acetate (3 x 25 mL). The combined organic extract was concentrated and purified by normal phase chromatography. The desired fractions were concentrated and the title compound was obtained as a white solid (1.6 g, 4.2 mmol, 62%). $^1$H NMR (300 MHz, Chloroform-*d*) δ 8.17 –

8.07 (m, 2H), 7.56 – 7.47 (m, 3H), 7.14 (s, 1H), 6.10 – 6.03 (m, 1H), 4.99 (s, 2H), 2.73 (d, *J* = 0.9 Hz, 3H), 2.37 (s, 3H), 1.50 (s, 9H).

**c) Preparation of 2-((2-(3,5-dimethyl-1*H*-pyrazol-1-yl)-6-phenylpyrimidin-4-yl)oxy)acetic acid**

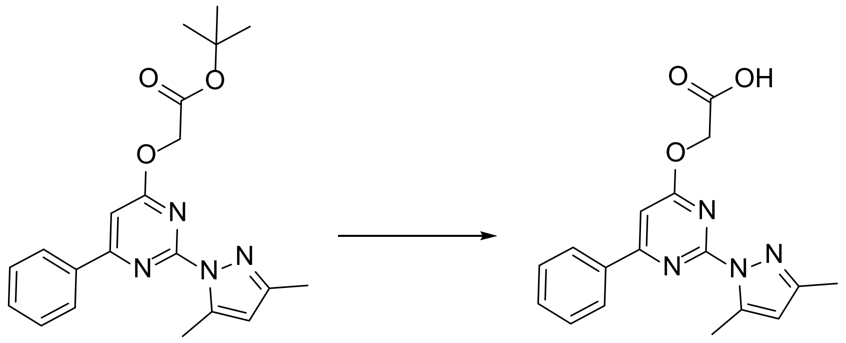

Trifluoroacetic acid (5 mL, 65 mmol) was added to a solution of *tert*-butyl 2-[2-(3,5-dimethylpyrazol-1-yl)-6-phenyl-pyrimidin-4-yl]oxyacetate (1.5 g, 3.9 mmol) in dichloromethane (5 mL). The mixture was stirred at room temperature for two hours. The reaction mixture was concentrated under vacuum. Ethyl acetate (20 mL) was added to the residue when a white solid crashed out. The solid was filtered, washed with ethyl acetate and dried at 60 °C when the title compound was obtained as a white solid (1.1 g, 3.4 mmol, 86%). $^{1}$H NMR (300 MHz, Chloroform-*d*) δ 8.11 – 8.00 (m, 2H), 7.58 – 7.47 (m, 3H), 7.17 (s, 1H), 6.08 (s, 1H), 5.47 (s, 2H), 2.82 (d, *J* = 0.8 Hz, 3H), 2.41 (s, 3H).

**d) Preparation of 2-[2-(3,5-dimethylpyrazol-1-yl)-6-oxo-4-phenyl-pyrimidin-1-yl]-*N*-methyl-*N*-(2-phenylethyl)acetamide hydrochloride.**

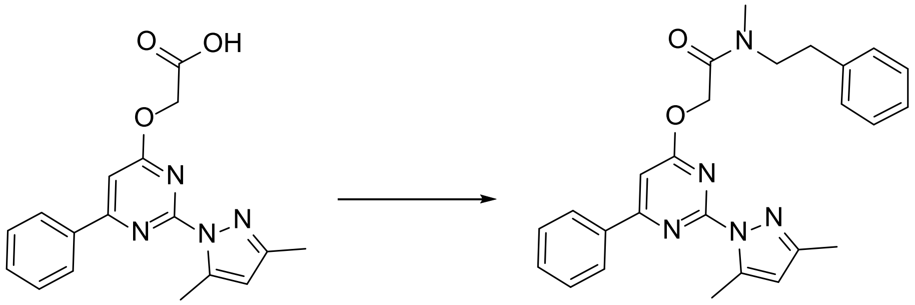

O-(7-Azabenzotriazol-l-yl)-*N,N,N',N'*-tetramethyluronium hexafluorophosphate (HATU, 0.11 g, 0.28 mmol) was added to a mixture of *N*-methyl-2-phenylethan-1-amine (58 mg, 0.43 mmol), 2-[2-(3,5-dimethylpyrazol-1-yl)-6-oxo-4-phenyl-pyrimidin-1-yl]acetic acid (70 mg, 0.22 mmol) and triethylamine (0.15 mL, 1.1 mmol) in dimethylformamide (1 mL) and the mixture was stirred overnight at ambient temperature. Water (10 mL) was added, and the product was extracted with ethyl acetate (3 x 20 mL). The combined organic extract was washed with brine and concentrated under vacuum. The crude product was purified by normal phase chromatography. The desired fractions were pooled and concentrated. The residue was dissolved in ethyl acetate (4 mL), and 2 M HCl in ether (2 mL) was added. The precipitated solid was isolated by centrifugation. $^{1}H$-NMR (300 MHz; $CD_3OD$) δ ppm (mixture of rotamers apparent): 8.32-8.25 (m, 2H), 7.63-7.49 (m, 4H), 7.40-7.25 (m, 5H), 6.71-6.68 (m, 1H), 5.36 and 4.70 (2s, 2H), 3.71-3.63 (m, 2H), 3.09 (s) and 3.05-2.95 (m) and 2.42-2.31 (m, 8H), 2.61-2.59 (2s, 3H); LRMS (ESI +): m/z 442.24 [$M+H^+$];

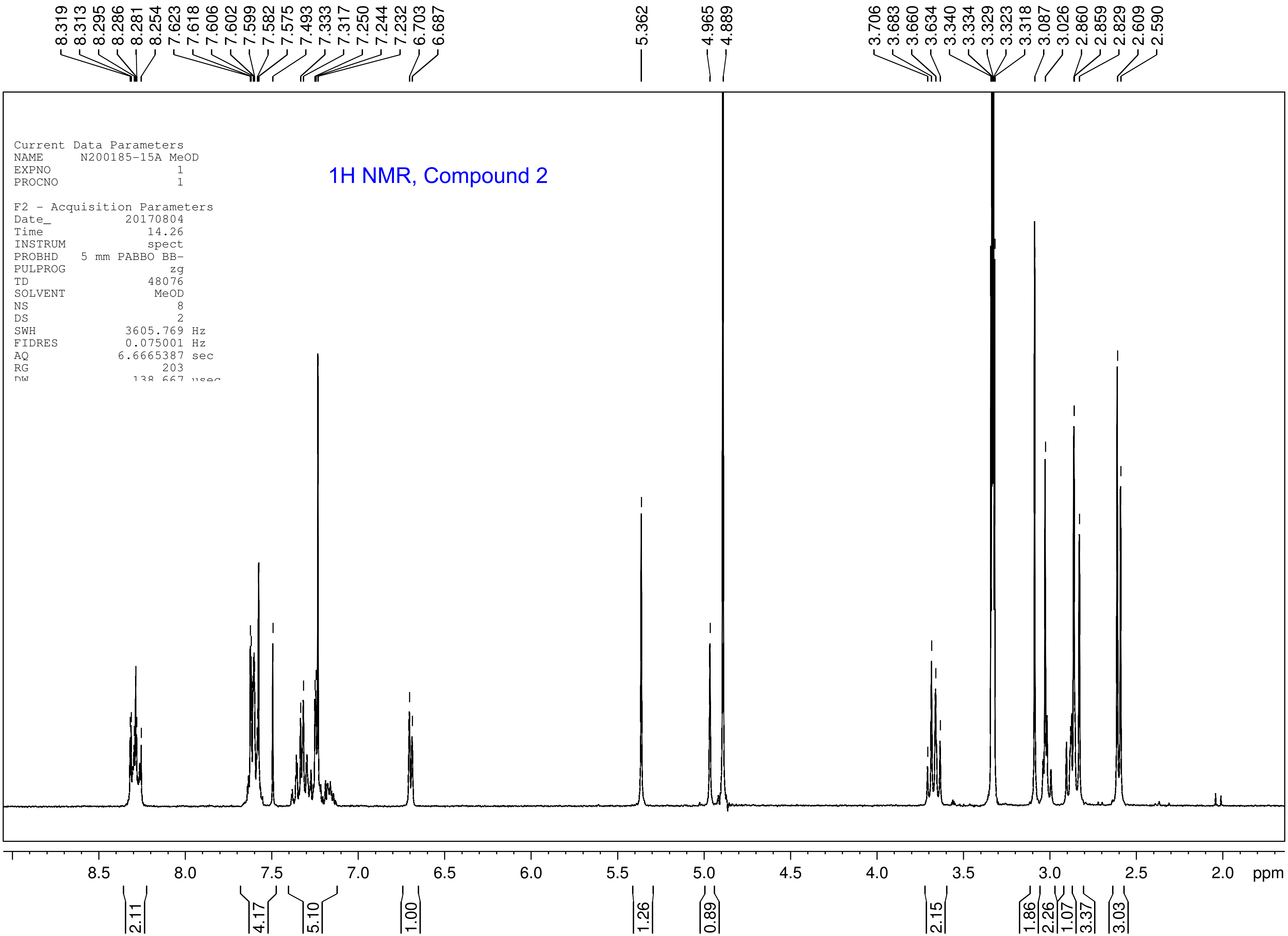
1H NMR, Compound 2
Current Data Parameters
NAME N200185-15A MeOD
EXPNO 1
PROCNO 1
F2 - Acquisition Parameters
Date_ 20170804
Time 14.26
INSTRUM spect
PROBHD 5 mm PABBO BB-
PULPROG zg
TD 48076
SOLVENT MeOD
NS 8
DS 2
SWH 3605.769 Hz
FIDRES 0.075001 Hz
AQ 6.6665387 sec
RG 203
8.319 8.313 8.295 8.286 8.281 8.254 7.623 7.618 7.606 7.602 7.599 7.582 7.575 7.493 7.333 7.317 7.250 7.244 7.232 6.703 6.687
5.362
4.965 4.889
3.706 3.683 3.660 3.634 3.340 3.334 3.329 3.323 3.318 3.087 3.026 2.860 2.859 2.829 2.609 2.590
8.5 8.0 7.5 7.0 6.5 6.0 5.5 5.0 4.5 4.0 3.5 3.0 2.5 2.0 ppm
2.11 4.17 5.10 1.00 1.26 0.89 2.15 1.86 2.26 1.07 3.37 3.03

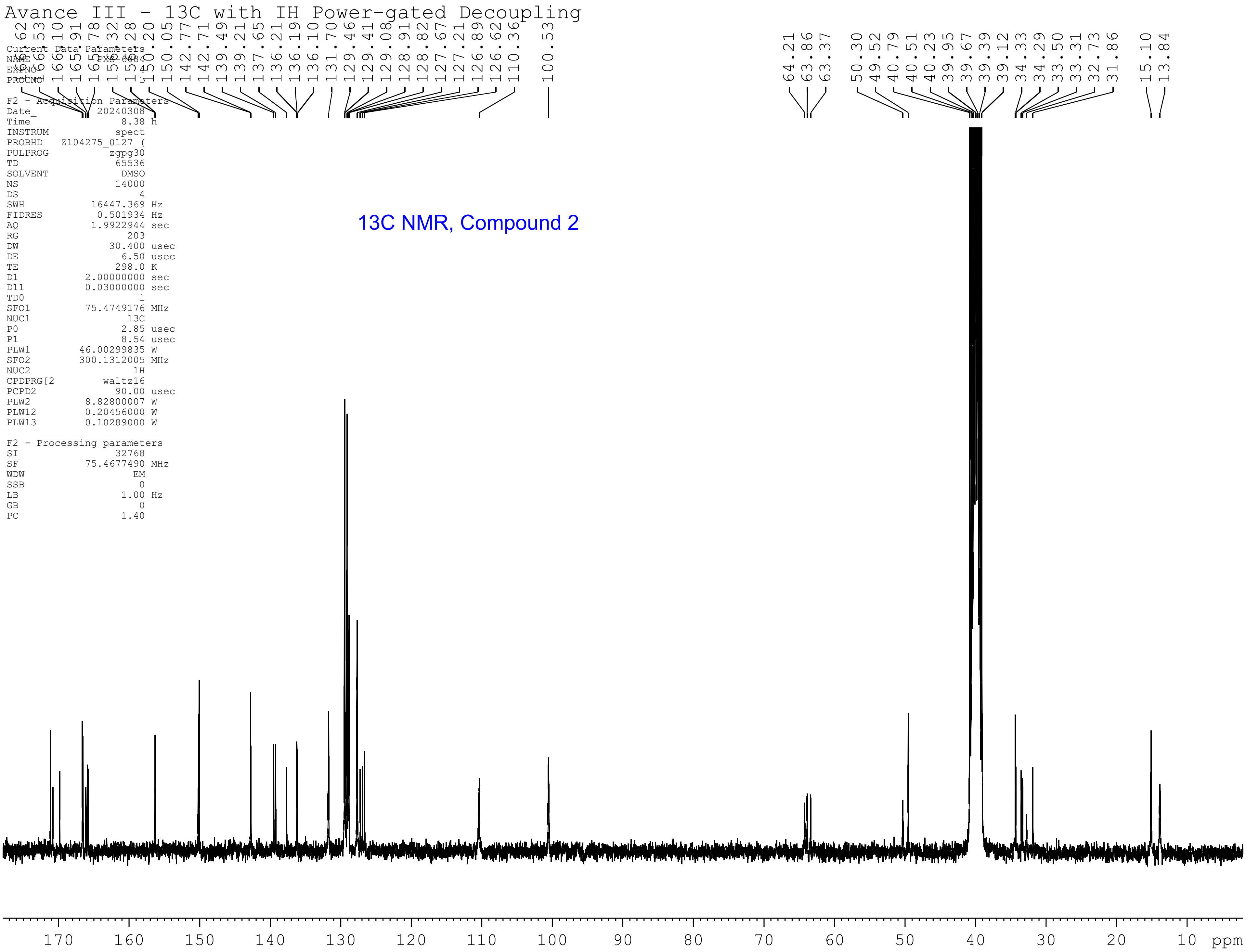
Avance III - 13C with IH Power-gated Decoupling
166.62
166.53
166.10
165.91
165.78
156.32
156.28
150.20
150.05
142.77
142.71
139.49
139.21
137.65
136.21
136.19
136.10
131.70
129.46
129.41
129.08
128.91
128.82
127.67
127.21
126.89
126.62
110.36
100.53
64.21
63.86
63.37
50.30
49.52
40.79
40.51
40.23
39.95
39.67
39.39
39.12
34.33
34.29
33.50
33.31
32.73
31.86
15.10
13.84
Current Data Parameters
NAME PXS-6084
EXPNO 4
PROCNO 1
F2 - Acquisition Parameters
Date_ 20240308
Time 8.38 h
INSTRUM spect
PROBHD Z104275_0127 (
PULPROG zgpg30
TD 65536
SOLVENT DMSO
NS 14000
DS 4
SWH 16447.369 Hz
FIDRES 0.501934 Hz
AQ 1.9922944 sec
RG 203
DW 30.400 usec
DE 6.50 usec
TE 298.0 K
D1 2.00000000 sec
D11 0.03000000 sec
TD0 1
SFO1 75.4749176 MHz
NUC1 13C
P0 2.85 usec
P1 8.54 usec
PLW1 46.00299835 W
SFO2 300.1312005 MHz
NUC2 1H
CPDPRG[2 waltz16
PCPD2 90.00 usec
PLW2 8.82800007 W
PLW12 0.20456000 W
PLW13 0.10289000 W
F2 - Processing parameters
SI 32768
SF 75.4677490 MHz
WDW EM
SSB 0
LB 1.00 Hz
GB 0
PC 1.40
13C NMR, Compound 2
170
160
150
140
130
120
110
100
90
80
70
60
50
40
30
20
10
ppm

HPLC, Compound 2

### LCMS Data - 254/280
### Chemistry Open Access

Sample Name : MD_N200185-15A
Sample ID :
Data Filename : MD_N200185-15AHPLV.lcd
Method Filename : QC_Everyday_Method_500ulpmin.lcm
Batch Filename : Chemistry Everyday Batch File.lcb
Vial # : 1-77
Sample Type : Unknown
Injection Volume : 5 uL
Date Acquired : 3/08/2017 5:29:18 PM
Acquired by : System Administrator
Date Processed : 3/08/2017 5:35:21 PM
Processed by : System Administrator

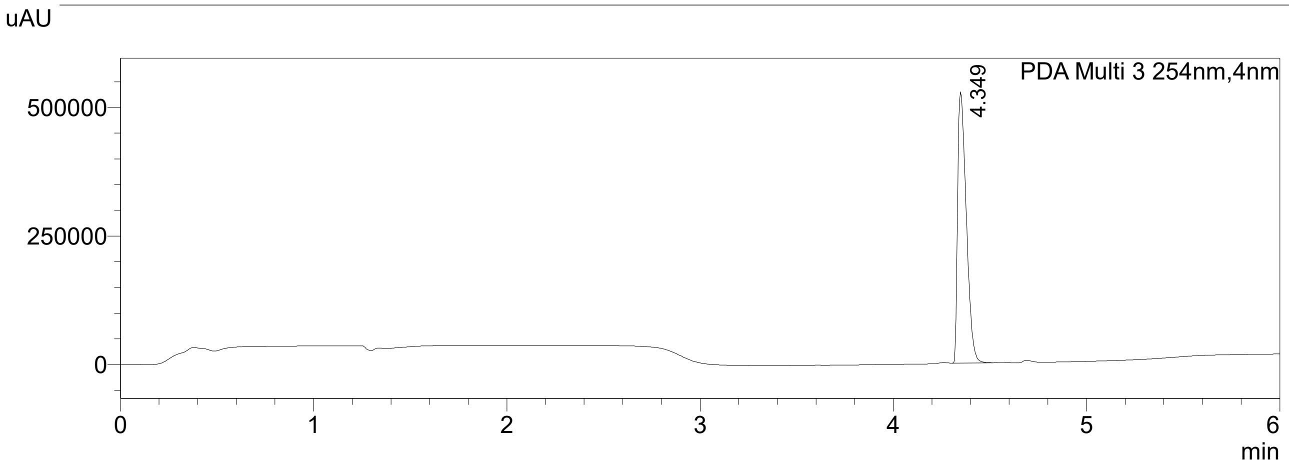


PDA Ch3 254nm

| Peak# | Ret. Time | Area | Area% |
|---|---|---|---|
| 1 | 4.349 | 1653486 | 100.000 |
| Total | | 1653486 | 100.000 |

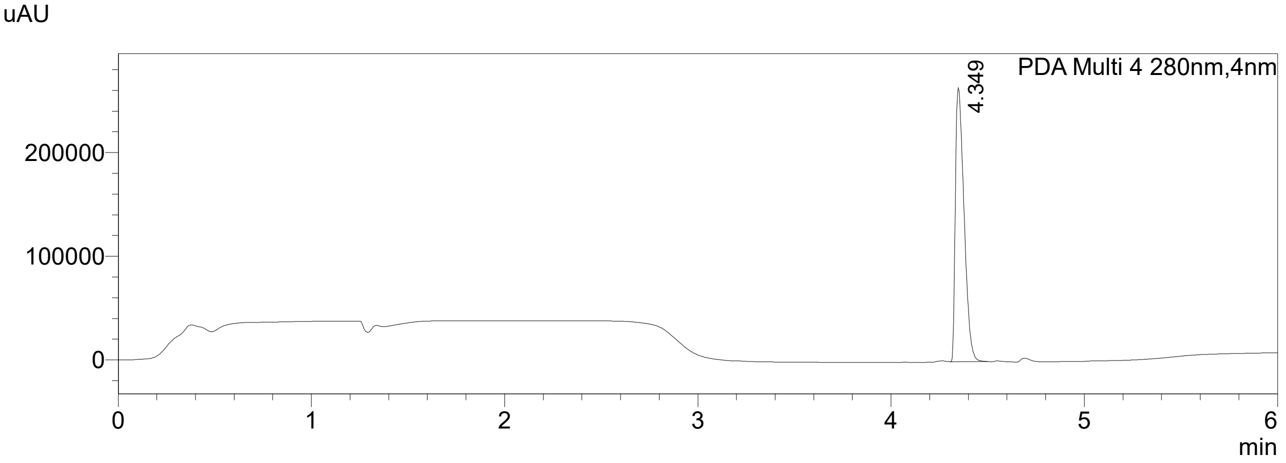


PDA Ch4 280nm

| Peak# | Ret. Time | Area | Area% |
|---|---|---|---|
| 1 | 4.349 | 828583 | 100.000 |
| Total | | 828583 | 100.000 |

Peak Profile

Peak# : 1
Retention Time : 4.349 min
Compound Name :
Wavelength Interval : 5nm

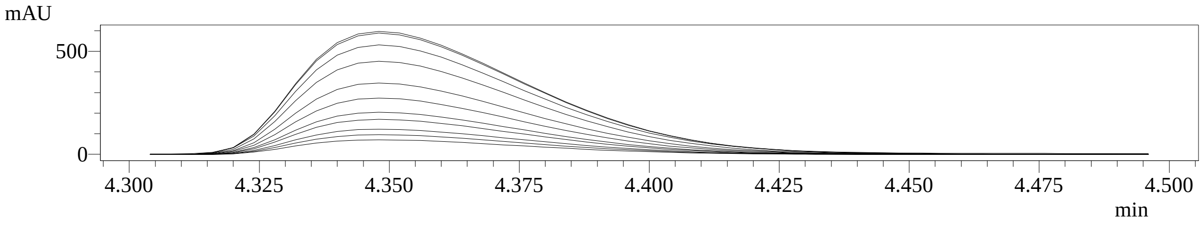

C:\Xcalibur\data\2017\JULY\MD_N200185-15A.raw 03/08/2017 16:23:10

Calibration Level
Client
Comment
Company
Dil Factor ..
ID ..
Injection Volume (µl) ..
Instrument Method ..
ISTD Amount ..
Laboratory
Phone
Processing Method ..
Row Number ..
Sample Type ..
Sample Volume (µl) ..
Sample Weight ..
Study
Vial ..

RT :0.00-6.00 SM: 3G

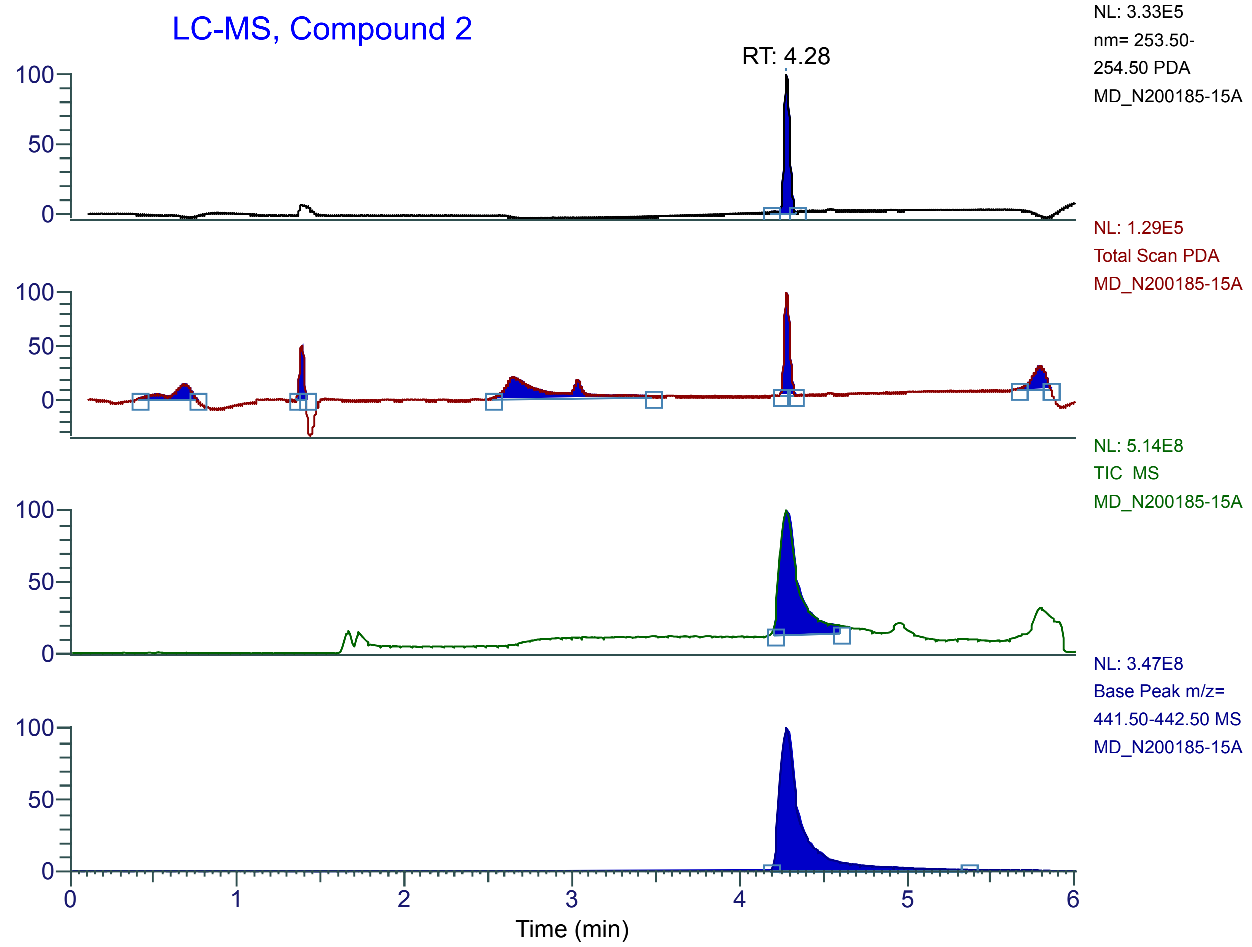


MD_N200185-15A #1251 RT: 4.28 AV: 1 NL: 3.97E5 microAU

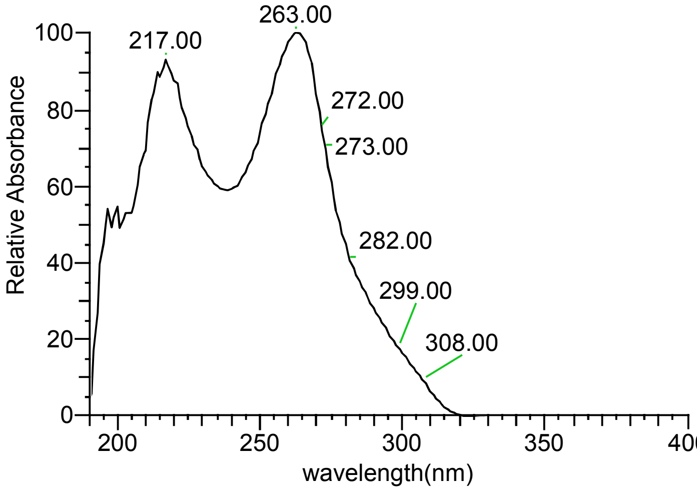


MD_N200185-15A #427 RT: 4.28 AV: 1 NL: 3.48E+008
T: + c ESI Q1MS [100.000-700.000]

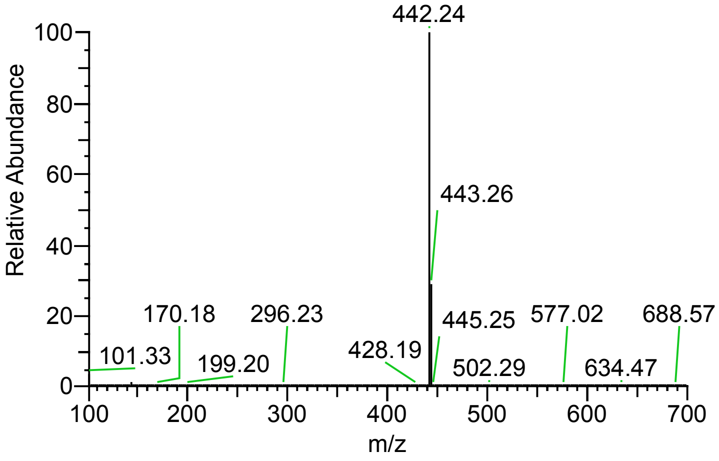


| RT (Min) | Peak Area | % Area |
|---|---|---|
| 0.686666666666667 | 154244.742697113 | 13.62 |
| 1.38666666666667 | 113804.184769496 | 10.05 |
| 2.65 | 425285.032453778 | 37.55 |
| 4.28333333333333 | 286536.520211035 | 25.3 |
| 5.79333333333333 | 152719.204143964 | 13.48 |

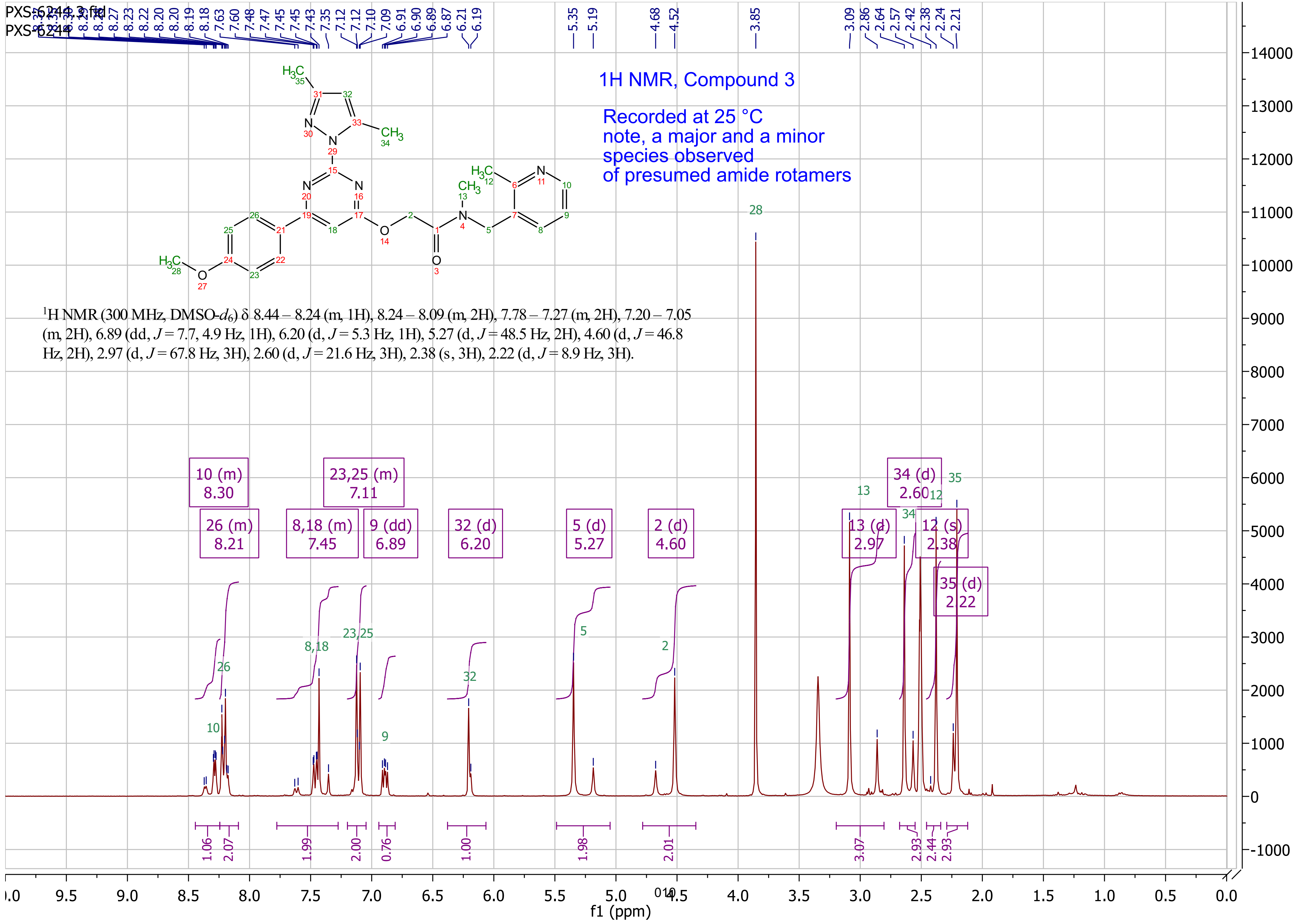

PXS-6244 3.fid
PXS-6244
1H NMR, Compound 3
Recorded at 25 °C
note, a major and a minor
species observed
of presumed amide rotamers
[1]H NMR (300 MHz, DMSO-$d_6$) δ 8.44 – 8.24 (m, 1H), 8.24 – 8.09 (m, 2H), 7.78 – 7.27 (m, 2H), 7.20 – 7.05 (m, 2H), 6.89 (dd, $J$ = 7.7, 4.9 Hz, 1H), 6.20 (d, $J$ = 5.3 Hz, 1H), 5.27 (d, $J$ = 48.5 Hz, 2H), 4.60 (d, $J$ = 46.8 Hz, 2H), 2.97 (d, $J$ = 67.8 Hz, 3H), 2.60 (d, $J$ = 21.6 Hz, 3H), 2.38 (s, 3H), 2.22 (d, $J$ = 8.9 Hz, 3H).
10 (m) 8.30
26 (m) 8.21
8,18 (m) 7.45
23,25 (m) 7.11
9 (dd) 6.89
32 (d) 6.20
5 (d) 5.27
2 (d) 4.60
13 (d) 2.97
34 (d) 2.60
12 (s) 2.38
35 (d) 2.22
1.06
2.07
1.99
2.00
0.76
1.00
1.98
2.01
3.07
2.93
2.44
2.93
f1 (ppm)

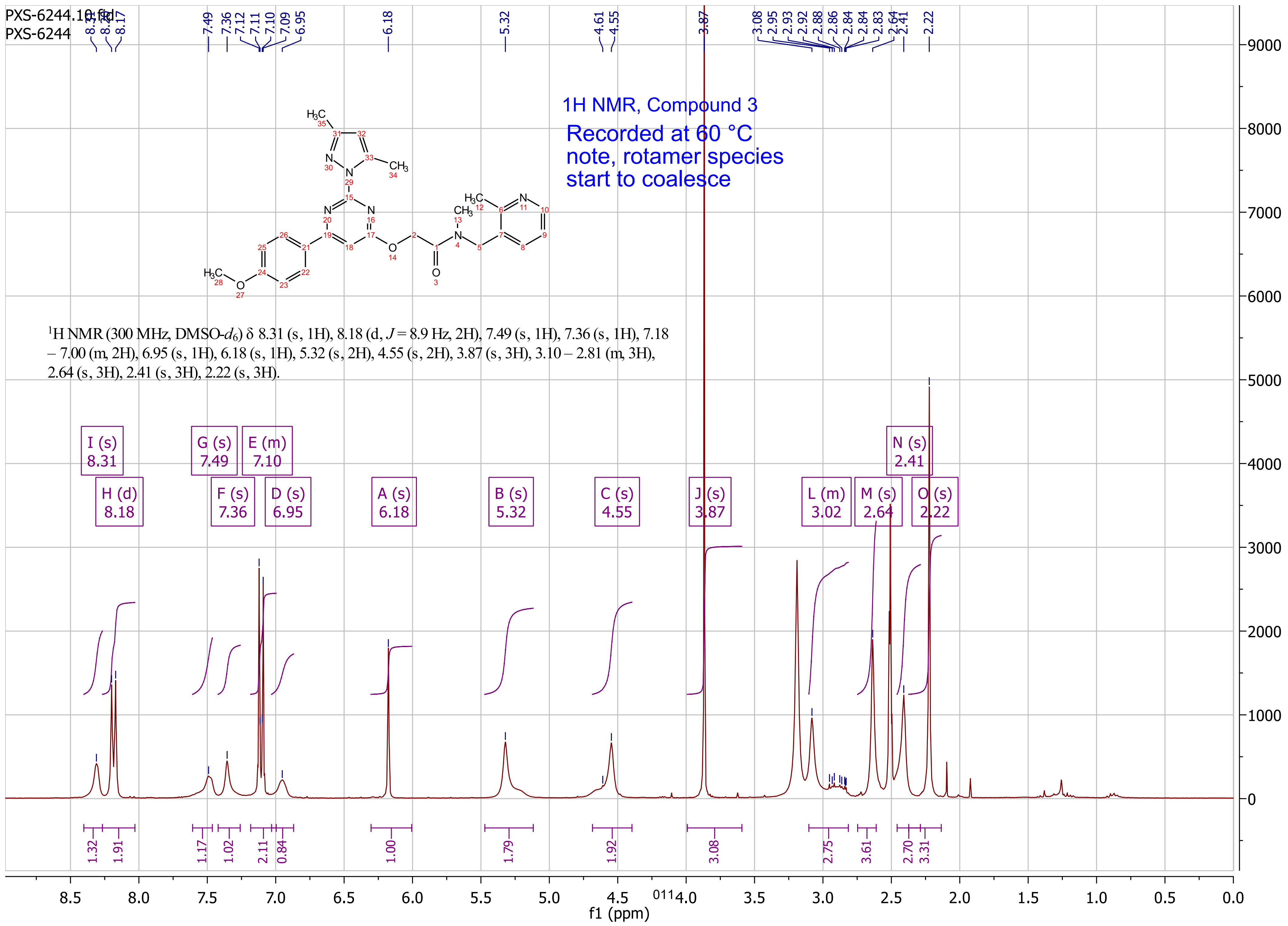
PXS-6244.10.fid
PXS-6244
1H NMR, Compound 3
Recorded at 60 °C
note, rotamer species
start to coalesce
$^{1}$H NMR (300 MHz, DMSO-$d_6$) δ 8.31 (s, 1H), 8.18 (d, J = 8.9 Hz, 2H), 7.49 (s, 1H), 7.36 (s, 1H), 7.18 – 7.00 (m, 2H), 6.95 (s, 1H), 6.18 (s, 1H), 5.32 (s, 2H), 4.55 (s, 2H), 3.87 (s, 3H), 3.10 – 2.81 (m, 3H), 2.64 (s, 3H), 2.41 (s, 3H), 2.22 (s, 3H).
f1 (ppm)

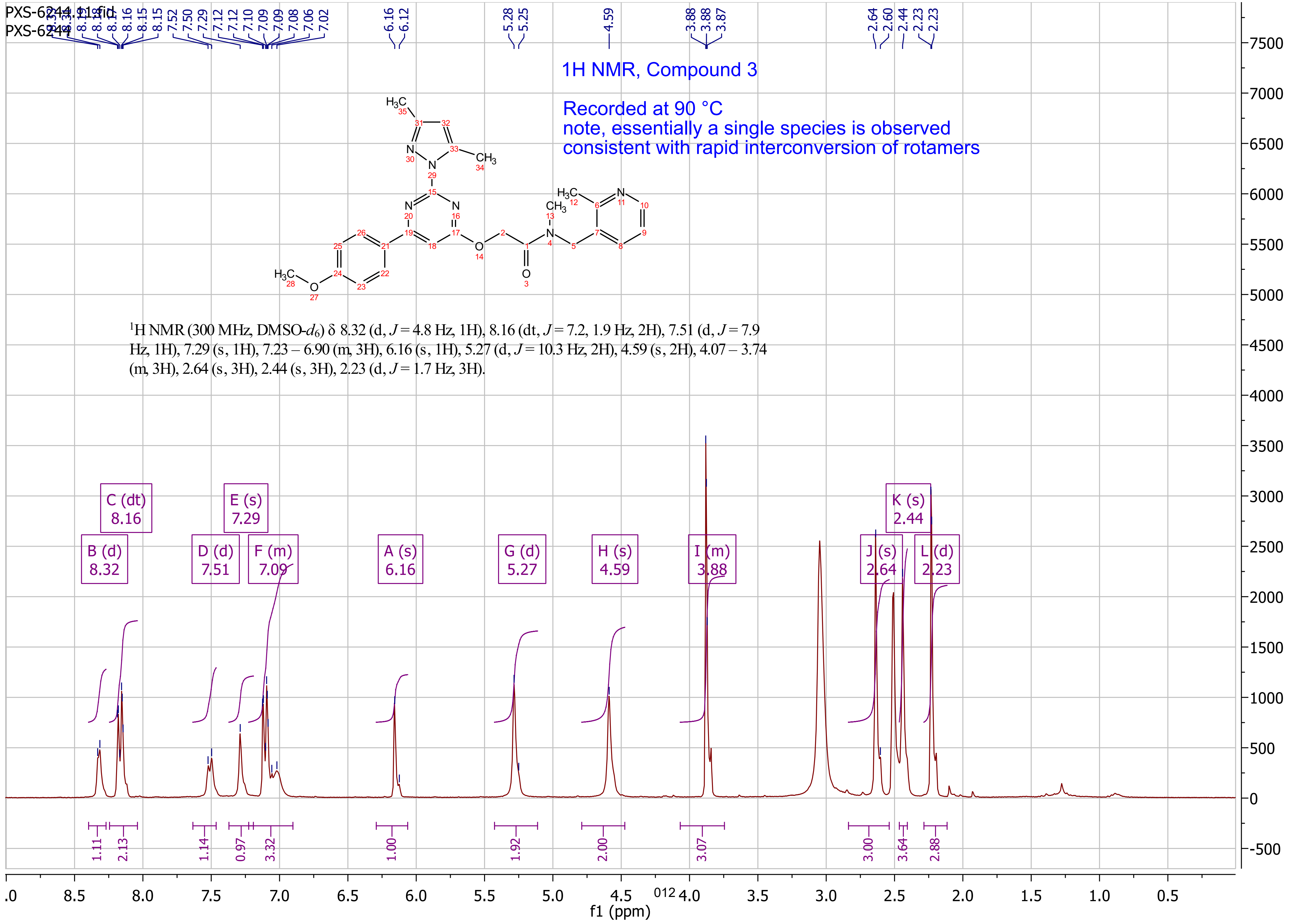
PXS-6244.11.fid
PXS-6244
1H NMR, Compound 3
Recorded at 90 °C
note, essentially a single species is observed
consistent with rapid interconversion of rotamers
[1]H NMR (300 MHz, DMSO-$d_6$) δ 8.32 (d, $J$ = 4.8 Hz, 1H), 8.16 (dt, $J$ = 7.2, 1.9 Hz, 2H), 7.51 (d, $J$ = 7.9 Hz, 1H), 7.29 (s, 1H), 7.23 – 6.90 (m, 3H), 6.16 (s, 1H), 5.27 (d, $J$ = 10.3 Hz, 2H), 4.59 (s, 2H), 4.07 – 3.74 (m, 3H), 2.64 (s, 3H), 2.44 (s, 3H), 2.23 (d, $J$ = 1.7 Hz, 3H).
B (d) 8.32
C (dt) 8.16
D (d) 7.51
E (s) 7.29
F (m) 7.09
A (s) 6.16
G (d) 5.27
H (s) 4.59
I (m) 3.88
J (s) 2.64
K (s) 2.44
L (d) 2.23
1.11
2.13
1.14
0.97
3.32
1.00
1.92
2.00
3.07
3.00
3.64
2.88
f1 (ppm)

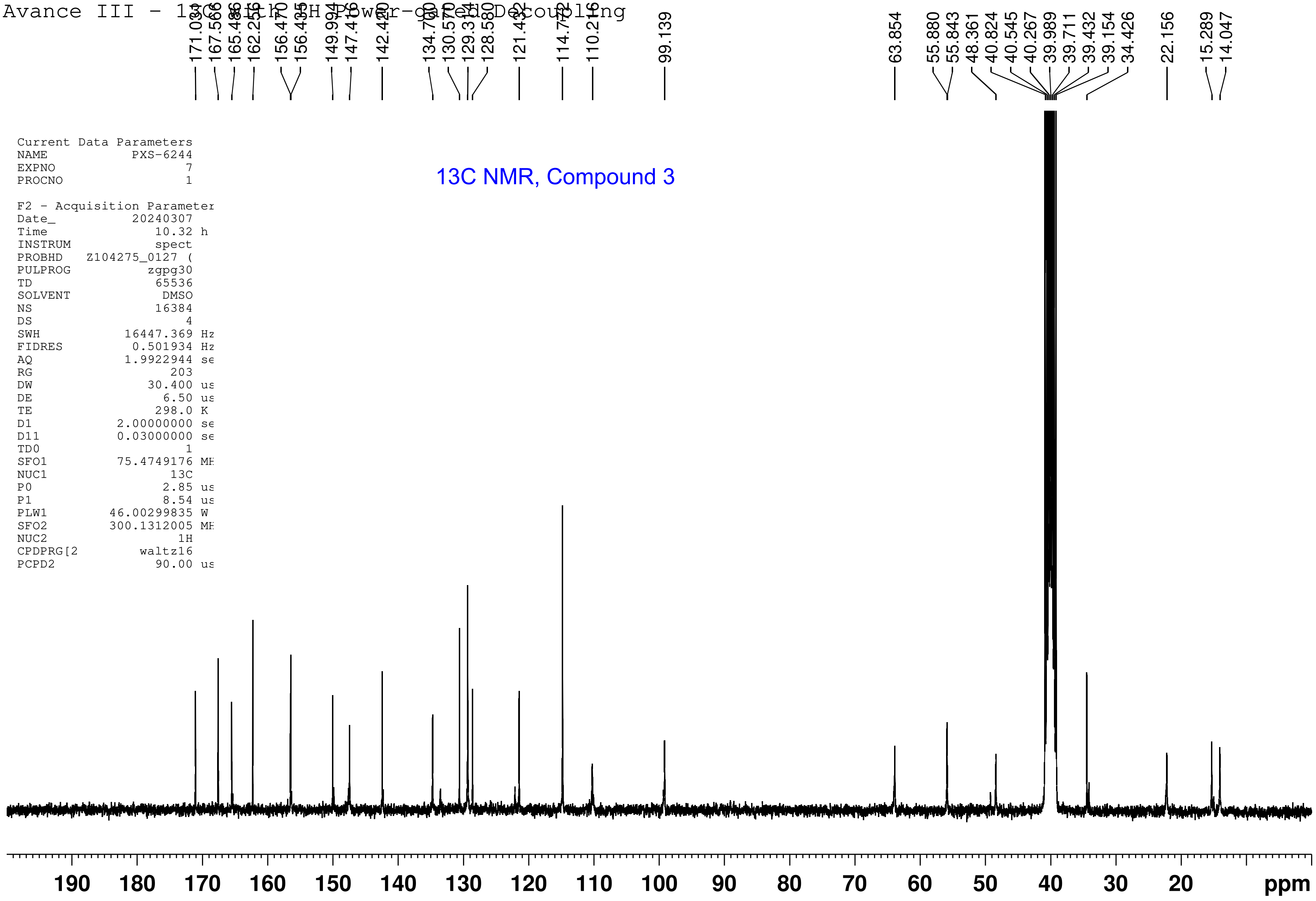
Avance III - 13C with 1H Power-gated Decoupling
171.030
167.566
165.486
162.255
156.470
156.435
149.934
147.416
142.420
134.740
130.570
129.314
128.580
121.432
114.772
110.216
99.139
63.854
55.880
55.843
48.361
40.824
40.545
40.267
39.989
39.711
39.432
39.154
34.426
22.156
15.289
14.047
Current Data Parameters
NAME PXS-6244
EXPNO 7
PROCNO 1
F2 - Acquisition Parameter
Date_ 20240307
Time 10.32 h
INSTRUM spect
PROBHD Z104275_0127 (
PULPROG zgpg30
TD 65536
SOLVENT DMSO
NS 16384
DS 4
SWH 16447.369 Hz
FIDRES 0.501934 Hz
AQ 1.9922944 se
RG 203
DW 30.400 us
DE 6.50 us
TE 298.0 K
D1 2.00000000 se
D11 0.03000000 se
TD0 1
SFO1 75.4749176 MH
NUC1 13C
P0 2.85 us
P1 8.54 us
PLW1 46.00299835 W
SFO2 300.1312005 MH
NUC2 1H
CPDPRG[2 waltz16
PCPD2 90.00 us
13C NMR, Compound 3
190 180 170 160 150 140 130 120 110 100 90 80 70 60 50 40 30 20 ppm

Sample Information

| | |
|---|---|
| Acquired by | : System Administrator |
| Date Acquired | : 4/03/2024 12:25:21 PM |
| Time Acquired | : 12:25:21 PM |
| Sample Name | : PXS-6244-04032024 |
| Tray# | : 1 |
| Vial# | : 84 |
| Injection Volume | : 5 |
| Data File | : 4032024_PXS-6244-04032024_01.lcd |
| Method File | : analytical_G3_10min_C3_1p5mlpmin.lcm |
| Background Data File | : |

LC-MS, compound 3

Chromatogram

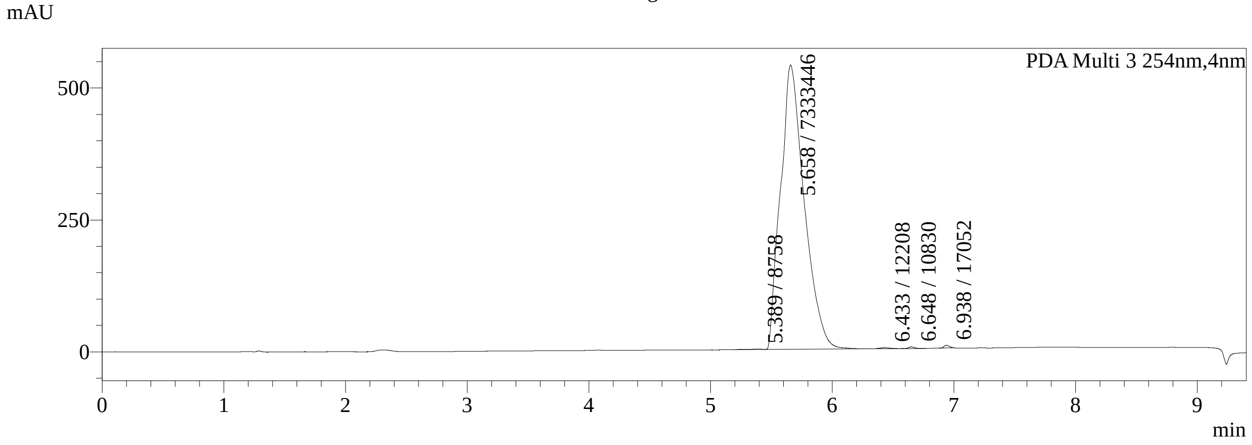


Peak Table

PDA Ch3 254nm

| Peak# | Ret. Time | Area | Area% |
|---|---|---|---|
| 1 | 5.389 | 8758 | 0.119 |
| 2 | 5.658 | 7333446 | 99.338 |
| 3 | 6.433 | 12208 | 0.165 |
| 4 | 6.648 | 10830 | 0.147 |
| 5 | 6.938 | 17052 | 0.231 |
| Total | | 7382293 | 100.000 |

Chromatogram

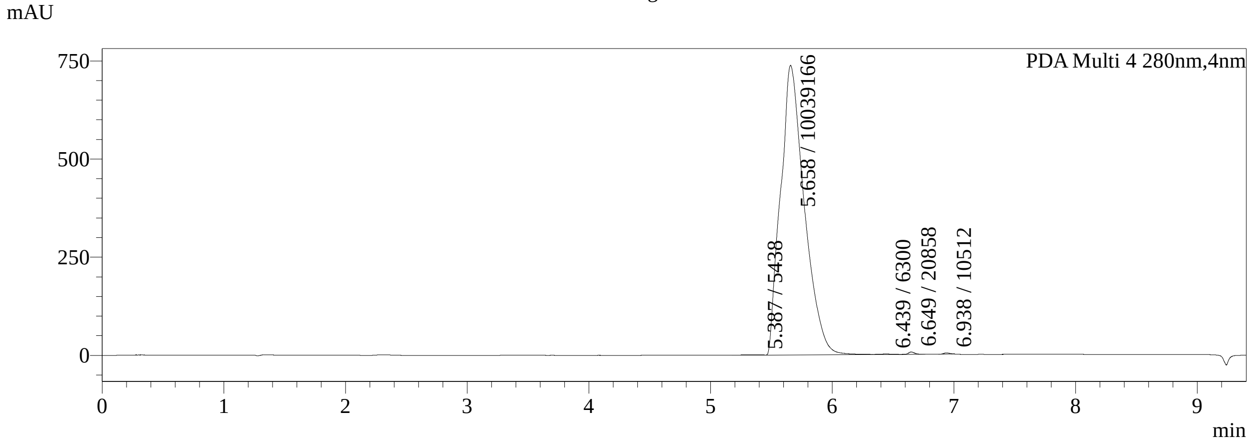


Peak Table

PDA Ch4 280nm

| Peak# | Ret. Time | Area | Area% |
|---|---|---|---|
| 1 | 5.387 | 5438 | 0.054 |
| 2 | 5.658 | 10039166 | 99.572 |
| 3 | 6.439 | 6300 | 0.062 |
| 4 | 6.649 | 20858 | 0.207 |
| 5 | 6.938 | 10512 | 0.104 |
| Total | | 10082274 | 100.000 |

Chromatogram

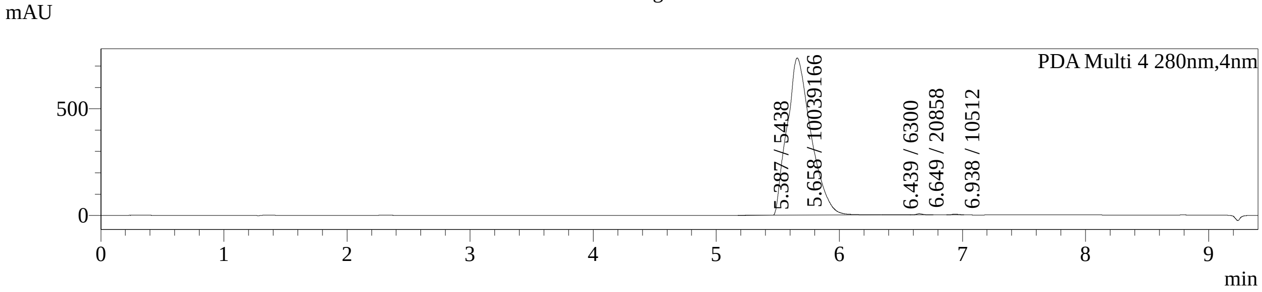


MS Chromatogram

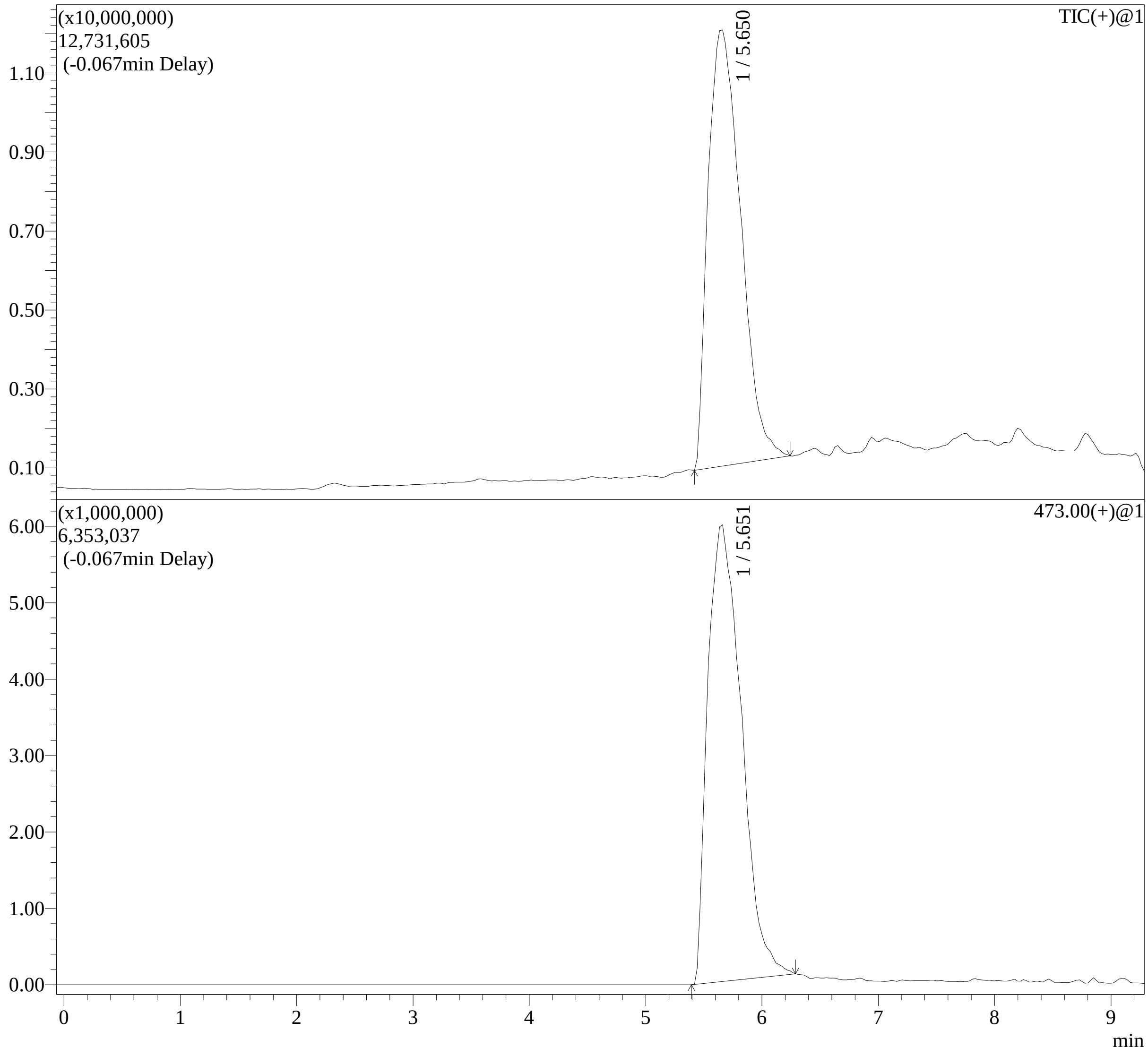

MS Spectrum

Line#:1 R.Time:----(Scan#:----)
MassPeaks:590
Spectrum Mode:Averaged 5.613-5.661(471-475) Base Peak:473(5835731)
BG Mode:Calc Segment 1 - Event 1

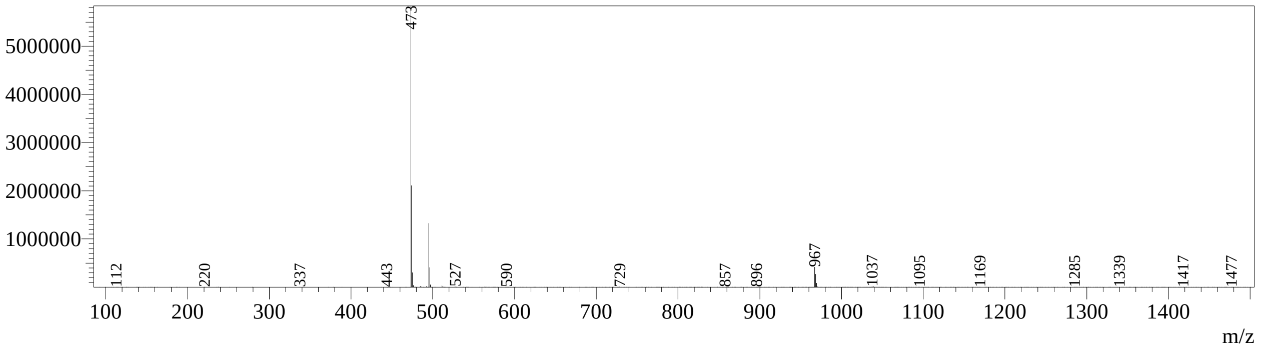


Line#:2 R.Time:----(Scan#:----)
MassPeaks:579
Spectrum Mode:Averaged 5.613-5.661(471-475) Base Peak:473(5850577)
BG Mode:Calc Segment 1 - Event 1

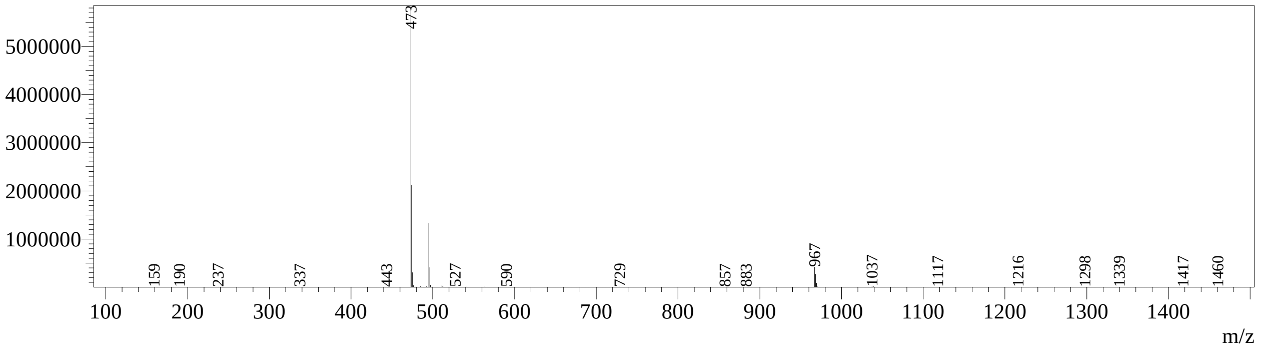

Method

<<Header>>

| | |
|---|---|
| Generated | : 26/11/2020 3:40:23 PM |
| GeneratedBy | : System Administrator |
| Modified | : 29/02/2024 5:35:00 PM |
| ModifiedBy | : System Administrator |

<<System Controller>>

| | |
|---|---|
| Model | : CBM-20A |
| Power On | : Off |
| Event1 | : Off |
| Event2 | : Off |
| Event3 | : Off |
| Event4 | : Off |
| Sample Load Timing | : Off |
| Sample Loading Overlap Time | : 0.00 min |

<<Data Acquisition>>

| | |
|---|---|
| LC Stop Time | : 9.40 min |
| AD2 Name | : MS Analogue |
| AD2 Sampling Frequency | : 2 Hz |
| AD2 Start Time | : 0.00 min |
| AD2 End Time | : 9.40 min |
| PDA Detector Name | : PDA |
| PDA Sampling Frequency | : 12.5 Hz |
| PDA Start Time | : 0.00 min |
| PDA End Time | : 9.40 min |
| PDA Time Constant | : 0.240 sec |

<<Pump>>

| | |
|---|---|
| Mode | : Binary gradient |
| Pump A | : LC-30AD |
| Pump B | : LC-30AD |
| Total Flow | : 1.5000 mL/min |
| B Conc. | : 5.0 % |
| B Curve | : 0 |
| PressMax | : 15000 psi |
| PressMin | : 0 psi |
| Solenoid Valve A Name | : Switching Valve |
| Solenoid Valve A | : A |
| Compressibility Setting | : On |
| Pump A Compressibility | : 0.45 /GPa |
| Pump B Compressibility | : 1.25 /GPa |

<<Autosampler>>

| | |
|---|---|
| Autosampler Model | : SIL-30AC |
| Enable Autosampler | : Use |
| Rinse Type | : External only |
| Sample Rack | : Rack 1.5 mL 105 vials |
| Rinsing Volume | : 500 uL |
| Needle Stroke | : 52 mm |
| Control Vial Needle Stroke | : 52 mm |
| Rinsing Speed | : 35 uL/sec |
| Sampling Speed | : 5.0 uL/sec |
| Measuring Line Purge Time | : 4.0 min |
| Rinse Port R0 Purge Time | : 4.0 min |
| Rinse Mode | : Before and after aspiration |
| Rinse Dip Time | : 3 sec |
| Cooler Temperature | : Off |
| Measuring Line Purge Volume | : 100 uL |
| Air Gap Volume | : Off |
| Rinse Method | : Rinse port only |
| Rinse Time | : 2 sec |

<<Oven>>

| | |
|---|---|
| Oven Model | : CTO-20AC |
| Enable Oven | : Use |
| Oven Temperature | : 40 C |
| Maximum Temperature | : 60 C |
| Rotary Valve L | : FCV-14AH |
| Rotary Valve R | : FCV-14AH |
| Valve L | : 3 |
| Valve R | : 3 |
| Ready Check | : Off |

<<LC Time Program>>

| Time | Module | Command | Value | Comment |
|---|---|---|---|---|
| 1.40 | Pumps | Pump B Conc. | 50 | |
| 6.00 | Pumps | Pump B Conc. | 95 | |
| 7.40 | Pumps | Pump B Conc. | 95 | |
| 7.41 | Pumps | Pump B Conc. | 5 | |
| 9.40 | Pumps | Pump B Conc. | 5 | |
| 9.40 | Controller | Stop | | |

<<Mobile Phase Name>>
Pump A Mobile Phase A : Water + FA
Pump B Mobile Phase A : Methanol

<<PDA>>
PDA Model : SPD-M20A
Lamp : D2&W

<<MS Parameter>>
Initial Valve Position :0
--Segment 1 Event 1--
Start Time :0.00 min
End Time :9.40 min
Acquisition Mode :Scan
Polarity :Positive
Event Time :0.45 sec
Detector Voltage :+1.00 kV
Threshold :100
Start m/z :90.00
End m/z :1500.00
Scan Speed :3750 u/sec
Interface Volt. :Use the Data in the Tuning File
DL Volt. :Use the Data in the Tuning File
Qarray DC Voltage :Use the Data in the Tuning File
Qarray DC Voltage :Use the Data in the Tuning File
--Segment 1 Event 2--
Start Time :0.00 min
End Time :9.40 min
Acquisition Mode :Scan
Polarity :Negative
Event Time :1.00 sec
Detector Voltage :+1.10 kV
Threshold :0
Start m/z :90.00
End m/z :1000.00
Scan Speed :938 u/sec
Interface Volt. :Use the Data in the Tuning File
DL Volt. :Use the Data in the Tuning File
Qarray DC Voltage :Use the Data in the Tuning File
Qarray DC Voltage :Use the Data in the Tuning File
<<MS Program>>

<<Adduct Ion>>
Adduct Ion :Use
--Positive--
+H :On
+Na :Off
+K :Off
+NH4 :Off
--Negative--
-H :On
+HCOO :Off
+CH3COO :Off
Charge :1

<<Peak Integration>>
<MS Analogue>
Channel : Ch1
Algorithm : Chromatopac
Width : 5 sec
Slope : 200 uV/min
Drift : 0 uV/min
T.DBL : 1000 min
Min.Area/Height : 1000 counts
Calculated by : Area
Auto : On
Auto Selection : Relative to Main Peak
Max. Peaks : 6
Relative to Main Peak : 10
Max Slices : 0
Peak Top Detection : Normal
RT Compensation Mode : Fine
Tailing Off : Off
Min.Area/Height is made effective in Manual Integration : Off
Noise Calculation Settings : Noise Data : Current Data
Calculation Method : ASTM
Range : Whole Range
Interval : 0.5 min
Include the Peak Detected Range : Off
Detection Limit Coefficient : 3.3
Quantitative Limit Coefficient : 10.0
Drift Calculation Settings : 0.000 - 15.000 min

<<Integration Time Program(Method)>>
<MS Analogue>
Channel : Ch1
Time Program : None

<<Integration Time Program(Data)>>
<MS Analogue>
Channel : Ch1
Time Program : None

<<Identification>>
<MS Analogue>
Window/Band : Window
Window : 5.00 %
Identification Method : Absolute
Peak Selection : Closest Peak
Display not identified peaks : Not display
Retention Time Update : None

<<Compound Table>>
<MS Analogue>
